# Inter-annual stability and age-dependent changes in plasma testosterone levels in a longitudinally monitored free-living passerine bird

**DOI:** 10.1101/2020.07.23.217836

**Authors:** Martin Tešický, Tereza Krajzingrová, Jiří Eliáš, Hana Velová, Jana Svobodová, Petra Bauerová, Tomáš Albrecht, Michal Vinkler

## Abstract

While seasonal trends in the testosterone-driven hormonal regulation of resource allocation are known from cohort population samples, data on the inter-annual individual stability of blood plasma testosterone levels in wild birds are lacking, and our understanding of age-dependent changes is limited. We assessed plasma testosterone levels in 105 samples originating from 49 repeatedly captured free-living great tits (*Parus major*) to investigate their relative long-term stability and lifetime changes. Furthermore, we examined the inter-annual stability of selected condition-related traits (carotenoid- and melanin-based plumage ornamentation, ptilochronological feather growth rate, body mass, and haematological heterophil/lymphocyte ratio) and their relationships to testosterone levels. We show that testosterone levels in both sexes are inter-annually repeatable, both in their absolute values and individual ranks (indicating the maintenance of relative status in a population), yet with higher stability in females. Despite this stability, in males we found a quadratic dependence of testosterone levels on age, with a peak in midlife. In contrast, female testosterone levels showed no lifetime trend. The inter-annual stability of condition-related traits was mostly moderate and unrelated to plasma testosterone concentrations. However, males with elevated testosterone had significantly higher carotenoid-pigmented yellow plumage brightness, presumably serving as a sexually selected trait. Showing inter-annual stability in testosterone levels, this research opens the way to further understanding of the causes of variation in fitness-related traits. Based on a unique longitudinal dataset, this study demonstrates that male plasma testosterone undergoes age-related changes that may regulate resource allocation. Our results thus demonstrate that male birds undergo hormonal senescence similar to mammals.

## Introduction

Testosterone (T) serves as a key endocrine mediator regulating the balance between investments into reproduction and various condition-related traits involved in self-maintenance (Fusani, 2008; Hau & Goymann, 2015; Kempenaers, Peters, & Foerster, 2008). Therefore, T levels may reflect individual condition, with links to immune function, physical activity, metabolism, growth, reproductive behaviours, and the expression of sexually selected ornamental traits (Hau and Goymann 2015). In bird populations, striking variations in T levels exist between different individuals, sexes, and throughout seasons. For example, in the blue tit (*Cyanistes caeruleus*) a ca. 200-fold difference exists between males with the highest and lowest T levels (Kempenaers et al., 2008). However, little is currently known about the individual inter-annual stability of T levels and T age-dependent trends in birds.

Avian T levels mirror many intrinsic and extrinsic factors (Kempenaers et al., 2008). Males have higher T levels than females, especially during breeding when T controls traits linked with mating, territorial aggressiveness, and parental care (Hau & Goymann, 2015). In temperate birds, male T levels peak with the increasing photoperiod during the early breeding season, followed by a decrease in the feeding period, after which T drops to steadily low levels (e.g. Van Duyse et al. 2003). Although less well understood, female plasma T follows a similar trajectory with lower amplitude and with a peak corresponding to the time of ovulation (Ketterson, Nolan, & Sandell, 2005). Importantly, variation in T levels reflects social status (Hau & Goymann, 2015), which is manifested through sexually selected condition-dependent carotenoid- (Albrecht et al. 2009; Svobodová et al. 2013) and melanin-pigmented ornaments (Griffith, Parker, & Olson, 2006; Guindre-Parker & Love, 2014).

Given the links between individual quality and T levels, long-term T stability in intra-population ranking can be predicted. A lack of T inter-annual stability would result in between-years variations in the related quality components. To our knowledge, studies specifically testing avian plasma T stability between years are presently lacking. Even in mammals, with more longitudinal data on repeatedly measured individuals (e.g. Perret 1992; Harman et al. 2001; Bernstein et al. 2012; Wolf et al. 2018), interannual T stability has only been rigorously tested in humans (Burger et al., 2000) and a few other mammals (Montano, Robeck, Steinman, & O’Brien, 2017; Perret, 1992).

After an initial increase of T during adolescence in both sexes, a higher amplitude of changes throughout ontogeny with a decrease during senescence is expected in males (Chahal & Drake, 2007). In mammals, this trajectory of hormonal senescence has been found in both cross-sectional and longitudinal studies, e.g. in humans (Harman et al., 2001; Vermeulen, Rubens, & Verdonck, 1972), macaques (*Ateles geoffroyi;* Hernández-López, Cerda-Molina, Díaz-Díaz, Chavira-Bolaños, & Mondragón-Ceballos, 2012) and beluga whales (*Delphinapterus leucas;* Montano et al., 2017). Although a similar pattern may be predicted also for birds, current evidence for age-related changes in avian T levels is mixed. A single study used longitudinal data which revealed a peak for middle-age birds and a subsequent age-related decline in T levels in the Florida scrub-jay (*Aphelocoma coerulescens;* Wilcoxen, Bridge, Boughton, Hahn, & Schoech, 2013). In all other (cohort) studies, age-related decreases in T levels have only been detected in the Japanese quail (*Coturnix japonica;* Balthazart et al. 1984), the common term (*Sterna hirundo;* Nisbet et al. 1999), the pied flycatcher (*Ficedula hypoleuca;* Moreno et al. 2014) and the barn swallow (*Hirundo rustica;* Adámková et al. 2019). In contrast, a majority of avian studies based on comparisons of young vs adult individuals (Belthoff et al. 1994; Schoech et al. 1996; Madsen et al. 2007) or cross-sectional data (Smith et al. 2005; Peters et al. 2002) reported no effect of age on male T levels. These results suggest inconsistency in age-dependent T changes in birds. Furthermore, little attention has been paid to female T levels despite their potentially non-negligible role in female physiology (Goymann & Wingfield, 2014) but see Moreno et al. (2014). Thus, longitudinal studies in free-living birds of both sexes investigating relationships between T age-related changes and fitness-related traits are lacking, but highly needed (Kempenaers et al., 2008).

In this article, we ask (1) whether plasma T levels in birds are inter-annually stable, and (2) if they undergo any ontogenetic changes. In a longitudinally monitored free-living great tit (*Parus major*) population, we obtained plasma samples and related data from 49 individuals of both sexes re-captured 2-3 times in subsequent years (2012-2017). Firstly, assuming links to individual quality, we predicted high inter-annual repeatability in individual T ranks, especially in males (Kempenaers et al., 2008). Secondly, assuming effects of ontogenetic development and senescence, we predicted age-dependent changes in plasma T levels with a maximum in midlife (Bouwhuis, Sheldon, Verhulst, & Charmantier, 2009). Thirdly, assuming life-history trade-offs mediated by T, we predicted negative associations between age-related T levels and condition-related traits (body mass and ptilochronologically measured feather growth rates, FGR), reflecting the organismal nutritional status (Grubb, 2006), but assumed a positive relationship between T levels and actual physiological stress (e.g. Deviche et al., 2014) assessed through haematological traits (heterophil: lymphocyte ratio, H/L ratio; Davis et al. 2008) on the one hand, and sexually selected ornaments (melanin-pigmented black breast stripe area and carotenoid-based plumage coloration (Hegyi, Szigeti, Torok, & Eens, 2007; Quesada & Senar, 2006) on the other. Finally, in T-dependent traits, we predicted inter-annual stability linked to T levels.

## Materials and methods

### Field procedures

A total of 886 adult free-living great tits were captured during breeding seasons (April–May) between 2012 and 2017 in a deciduous forest at the edge of Prague, Czech Republic (for a detailed description of the study locality and field procedures, see Supplementary materials and methods online 1; SMMO 1 in SI), in total 49 individuals (28 males and 21 females) and repeatedly sampled for blood plasma (for age-structure see Fig. S1). The study area is located near the Kbely meteorological station (Czech Hydrometeorological Institute), where information on mean air temperature seven days before capturing (further referred to as temperature) was obtained. To standardise sampling, all birds were captured into a mist net when their nestlings were 7-14 days old. Immediately after capture, approximately 150 μl of blood was collected from each individual by an insulin syringe (Omican 50-50I.U./0,5ML 30G x12; B. Braun, Melsungen, Germany) from a jugular vein. Two blood smears per individual were prepared from drops of blood, part of the sample was stored frozen in a microtube with 96% ethanol for later use (not used in this study) and a subsample of the blood (ca. 70-100 μl) was stored in a cool box and transported later the same day into the laboratory. Each blood sample was centrifuged in a microcentrifuge (type 5424, Eppendorf, Hamburg, Germany) at 8000 rpm for 5 minutes and the separated plasma was frozen at −80 °C. In the field, weight (measured by a digital scale, *d* = 0.02 g, type PPS200, Pesola, Schindellegi. Switzerland) and tarsus length (measured by a digital calliper, accuracy 0.01 mm; Kinex, Prague, Czech Republic) were recorded to later calculate the size-standardised body mass as the ratio of weight to tarsus length (further referred to as body mass). The second outermost tail rectrix from the left side was collected from each individual for the ptilochronological assessment (Grubb, 2006). To assess melanin-based and yellow carotenoid-based plumage ornamentation, we first collected a standardised digital image of the black breast stripe using a scanner (type Perfection V30, Seiko Epson, Nagano, Japan; Bauerová et al., 2017). Then, samples of ornamental feathers from the upper part of the yellow breast area were collected for later light spectral analysis (ca. 20-25 feathers). Finally, each individual was tagged with a steel ring with a unique code of the Czech Bird Ringing Centre, National Museum in Prague. The minimum age of each bird was estimated based on a combination of the ringing records and plumage age characteristics at their first capture (Svensson & Baker, 1992).

### Plasma testosterone assay

Plasma T was quantified in duplicates using a Testosterone ELISA Kit (480 solid well; product No 582701; Cayman Chemical Company, Ann Arbor, USA) according to the manufacturer’s instructions, with measuring T range 3.9-500 pg/ml and sensitivity 6 pg/ml, using 20 μl of the plasma sample diluted 5.5× as input. The measurement repeatability in duplicates was *r* = 0.91 (*p* < 0.001) and intra- and inter-assay variation coefficients (CV) were 13.01 (N =102) and 24.01 % (N = 6), respectively (see SMMO 2 in SI for details).

### Condition-related traits

For the haematological analysis we followed the procedure reported previously by Bauerová et al. (2020). From blood smears stained with Wright-Giemsa Modified stain (product No. WG128, Sigma-Aldrich, St. Luis, MO, USA), we calculated the differential white blood cell count using a light microscope with 100× objective magnification (Olympus Corporation, Tokyo, Japan, type CX-31), which served to estimate the H/L ratio (see SMMO 3 in SI for details).

To evaluate the individual nutritional status in the moulting period, we performed a ptilochronological analysis using the previously collected rectrix samples (Vinkler, Schnitzer, Munclinger, & Albrecht, 2012). After scanning all rectrices (scanner Epson V30) and post-processing the images in Corel Photo-Paint X3 software (Corel Corporation, Ottawa, Canada), we estimated the FGR as the mean growth bar width in a segment of 10 growth bars with the centre located at 2/3 of the feather length using ImageJ software (v. 2.0.0; Schindelin et al. 2015; see SMMO 4 in SI for details).

From the scanned digital images of the black breast stripe ornaments, the breast stripe area was measured in Adobe PHOTOSHOP CS. 2 software (v. 10.0; Adobe Systems, San Jose, USA) according to Bauerová et al. (2017). For 20 randomly selected individuals, the repeatability of the measurement was found to be high (*r* = 0.9, *p* < 0.001).

Yellow carotenoid-based breast ornament was analysed using an Avaspec 2048 spectrometer with an Avalight XE light source and the Avasoft 7.0 processing system (Avantes, Eerbeek, Netherlands). For each individual, we measured the colour of a layer of 20-25 carotenoid-based feathers fixed on a glass slide (according to Quesada and Senar 2006). As estimates of the yellow colouration parameters, we calculated the chroma (the difference between the reflectance at 700 nm and 450 nm, relative to the reflectance at 700 nm; the interval for absorbance of carotenoid pigments is 450–700 nm) and total brightness (total reflectance, i.e. the sum of reflectance in visible light from 300 to 700 nm) from the spectral data (Montgomerie, 2006; see SMMO 5 in SI and Svobodová et al. 2018 and Albrecht et al. 2009 for more details).

### Statistical analyses

Statistical analyses were performed using R software (v. 3.4.1; R Core Team, 2017). First, to control for technical factors possibly affecting plasma T measurements (i.e. those that were not biologically relevant), we built a Linear Model (LM) with log-transformed T (T_raw_) as the dependent variable, and testosterone plate (batch in which the sample was measured), time of capture and handling time (i.e. the time between capture and blood sampling) as explanatory variables (model M1, Table S3 in SI). We then used the residuals for testosterone from this model (termed M1) as ‘controlled T values’ (further referred to as T) in all subsequent analyses.

We checked the inter-annual stability of T and condition-related traits separately for both sexes using two complementary calculations of repeatability, both based on variance components from Linear Mixed Models (LMMs) with individual ID as a random factor using the rptrR package, version 9.22 (Stoffel, Nakagawa, & Schielzeth, 2017). First, we assessed the inter-annual repeatability of absolute trait values (further referred to as absolute repeatability), and second, the inter-annual repeatability of the weighted relative rank of the trait in the population (i.e. relative order for a given trait weighted by the number of observations in each year; further referred to as individual rank consistency). Some traits can function as relative traits that possess low inter-annual repeatability in absolute values, but their relative status (rank) is maintained in a population between years (Senar & Quesada, 2006).

To see whether T levels undergo similar trajectories over the lifetime of both sexes, we tested for sex-specific age-related changes in plasma T levels in an LMM model (M2) with T levels used as a dependent variable, and sex, age (also as a quadratic term), their interactions (sex:age, sex:age ^2), Julian date (the date of capture converted to the Julian calendar), mean air temperature (mean air temperature seven days before catching; SMMO 1), tarsus length and selected condition-related traits (mass, FGR, H/L ratio and H/L ratio:sex) as fixed effects. The year of capture and individual ID were used as variables with random intercept effects, while the age was given a random slope effect within the individual ID (age|ID) to allow for inter-individual variations in age-related changes.

To investigate relationships between plasma T levels and ornament expression, we built LMM models separately for both sexes with the same random factors as in M2 and breast stripe area (M3-4), yellow chroma (M5-6) and yellow brightness (M7-8) used as dependent variables. In these models, the explanatory variables used as fixed effects were age, T, tarsus length, Julian date, and condition-related traits (mass, FGR, H/L ratio).

To achieve normality in model residuals (checked using the Shapiro-Wilk normality test), some response variables were log-transformed before model testing (Table S3 in SI). Minimum adequate models (MAMs; defined here as models with all fixed terms significant at the level of p ≤ 0.05 or with marginally insignificant terms at the level of p < 0.10) were selected by backward elimination of non-significant terms from the full models (Crawley, 2013). The backward elimination steps in the models were checked by changes of deviance with accompanied changes in degrees of freedom (ANOVA) and Akaike information criterion (AIC) using F statistics. For LMMs models, the packages nlme; version 3.1 (Pinheiro, Bates, DebRoy, & Sarkar, 2019) and lmer4; versions 1.1 (Bates, Mächler, Bolker, Benjamin, & Walker, Steven, 2015) were used.

## Results

### Inter-annual stability and individual rank consistency in T levels

In our set of 105 samples originating from 49 re-captured individuals, mean plasma T_raw_ concentrations were 393.93 ± 388.24 pg/ml in males (N = 28) and 110.43 ±74.78 pg/ml in females (N = 21; see Table S1 for plasma T_raw_ concentrations across the age cohorts). Given the differences in T levels between the sexes (unpaired two-sample t-test; *t* = 7.185, *df* = 91.685, *p* < 0.001), we calculated their inter-annual T stability separately. The absolute inter-annual repeatability was lower for males (*r* = 0.103; *p* > 0.05) than for females (*r* = 0.576; *p* < 0.001) (Fig. 1A; Table S2A and Fig. S2A-B in SI), but not considerably different when relative ranks were taken into account (in males *r* = 0.411; *p* = 0.008; in females *r = 0.489; p = 0.005;* Fig. 1B; Table S2B in SI). Absolute T levels of two-year-old individuals did not correspond well with their levels in subsequent years, because excluding observations of the two-year-old individuals from the data set considerably increased the absolute T inter-annual repeatability in males (*r* = 0.385; *p* = 0.045), but not in females (*r* = 0.59; *p* = 0.016; Table S2A).

**Figure 1.**
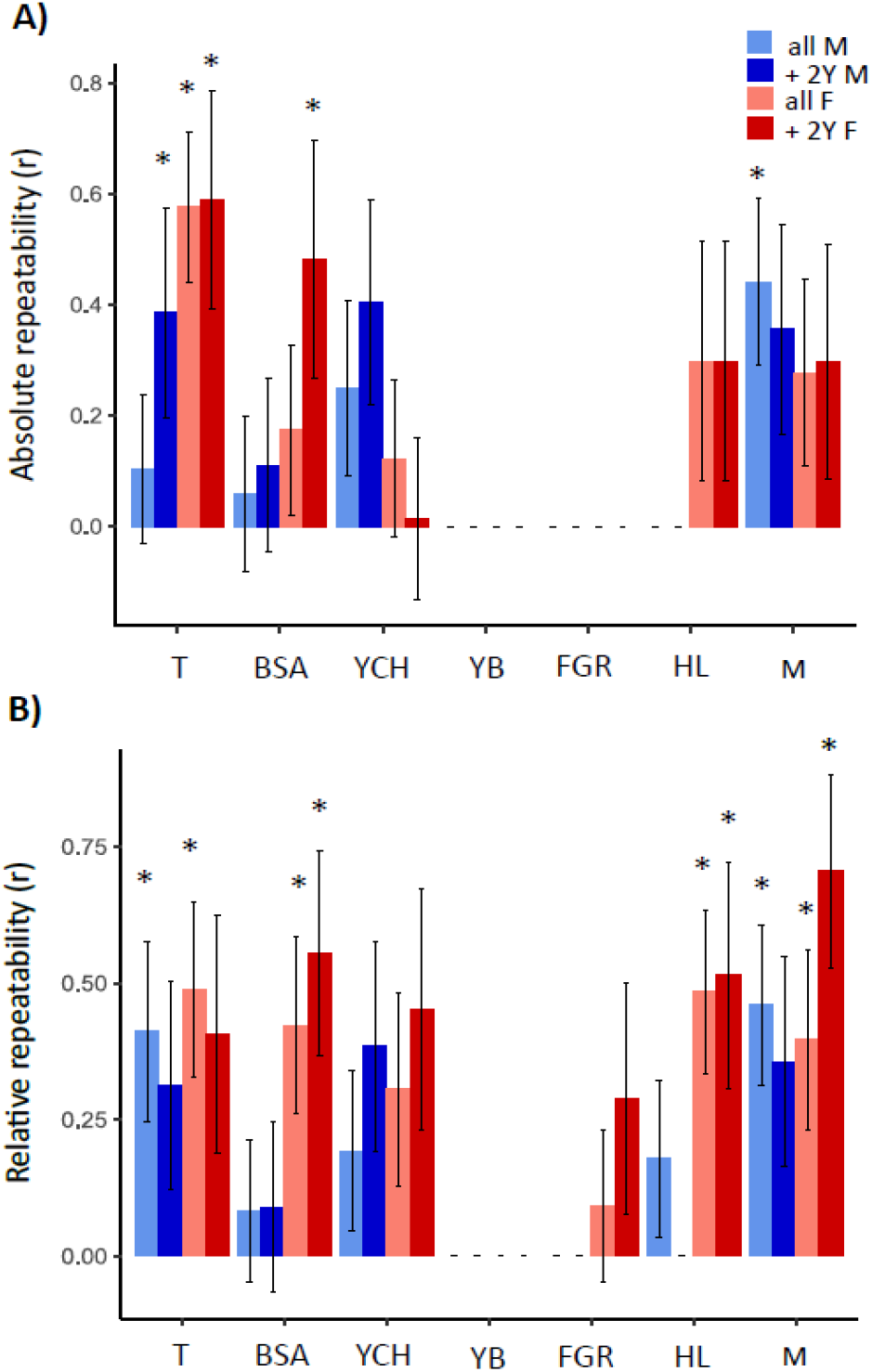
Inter-annual individual absolute (A) and relative (B; also termed individual rank consistency) repeatability of testosterone and condition-related traits in repeatedly captured great tits. The repeatability is indicated by the bar height with the standard error, separately for each category (for cases with no repeatability no bars are shown). *P*-values at statistical significance level *p* ≤ 0.05 are marked with an asterisk (see Table S2 in SI for details). T – residuals for plasma testosterone, BSA – the black breast stripe area, YCH – chroma of yellow ornamental plumage, YB – brightness of yellow ornamental plumage, FGR – feather growth rate, i.e. daily growth bar width increase in rectrices, HL – the ratio of heterophils to lymphocytes in peripheral blood, M – body mass (weight relative to tarsus length). The calculations were done separately for both sexes and with either all observations or after excluding observations of young birds (two-year-olds; 2Y): All M – all male individuals (N_md_ = 28, N_obs_ = 58), +2Y M – only male individuals older than two years (N_md_ = 17, N_obs_ = 36), all F – all female individuals (N_ind_ = 21, N_obs_ = 47), +2Y F – only female individuals older than two years (N_ind_ = 11, N_obs_ = 24)

### Age-dependent changes in plasma T levels

Next, we tested whether T levels change during the lifetime (MAM2; Table 1, Table S3 in SI). We found a significant effect of age (age: slope = 0.032; *p* = 0.003 and age^2: slope = 0.462; *p* = 0.059) on T levels with sex-specific interactions (age^2:sex slope = −3.390, *p* = 0.032 and age:sex slope = −2.887, *p* = 0.005), with male T concentrations peaking in middle-age and decreases in older birds, while in females levels were stable throughout life (Fig. 2).

**Figure 2.**
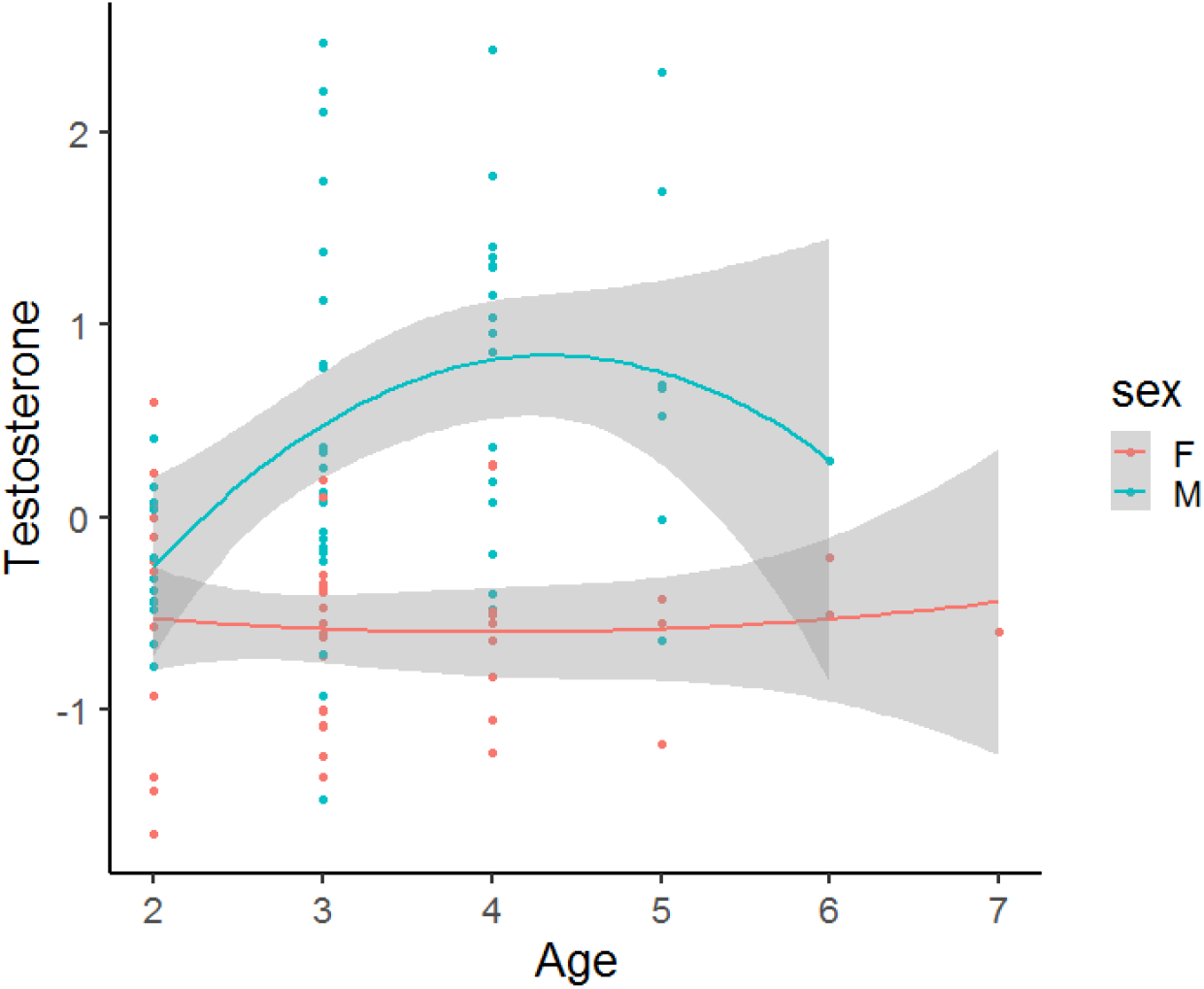
The relationship between plasma testosterone (T) levels and age in re-captured great tit males and females (N_md_ = 49, N_obs_ = 105). Plasma T levels are shown from minimal adequate model 2 (Table 1), with confidence intervals indicated by grey shaded areas. Age is the minimal estimated age based on bird ringing

**Table 1.** Minimum adequate models (MAMs) for re-captured great tits. For Linear Mixed Models (MAM2 – 8), the year of capture and individual ID were used as random effects with random intercepts. Age was given as a random slope effect within the individual ID (age|ID). Residuals for testosterone (T) from full Linear Model 1 were used in all other models. Slope ± SE values are provided only for continuous variables. *P*-values at statistical significance level *p* ≤ 0.05 are in bold. T – plasma testosterone concentration [pg/ml], breast stripe – the black breast stripe area [mm^3^], individual – individual identification based on bird ringing, sex, age – minimal estimated age based on bird ringing (age^2 – polynomic term), T_ave7 – mean daily temperature 7 days before catching, FGR – feather growth rate, Jul.date – Julian date from the beginning of the year, mass – body mass as the ratio of weight and tarsus length, HL – HL ratio (number of heterophils/number of lymphocytes), brightness – brightness of yellow carotenoid ornament, yellow chroma –chroma of the yellow carotenoid ornament, N_obs_ – number of observations, N_md_ – number of individuals

| Minimum adequate model | Slope ± SE | F | Df | p | N<sub>obs</sub>/N<sub>ind</sub> |
| --- | --- | --- | --- | --- | --- |
| MAM 1 $\log(T_{raw}) \sim T_{plate}$ | | 2.766 | 5/104 | 0.022 | 105/49 |
| MAM 2 $T \sim \text{sex} + \text{age} + \text{age}^2 + T_{ave7} + \text{sex}:\text{age}^2 + \text{sex}:\text{age} + (1\vert \text{year}) + (\text{age}\vert \text{individual})$ | | 43.005 | 6/6 | <0.001 | 105/49 |
| sex | 0.990 ± 0.172 | 31.408 | 3/9 | <0.001 | |
| age | 0.032 ± 0.100 | 15.870 | 4/8 | 0.003 | |
| age<sup>2</sup> | 0.462 ± 1.029 | 5.665 | 2/10 | 0.059 | |
| T<sub>ave7</sub> | -0.056 ± 0.031 | 3.248 | 1/11 | 0.071 | |
| sex:age<sup>2</sup> | -3.390 ± 1.633 | 4.589 | 1/11 | 0.032 | |
| sex:age | 2.877 ± 1.690 | 10.459 | 2/10 | 0.005 | |
| MAM 3 (males): $\log(\text{breast stripe}) \sim \text{FGR} + (1\vert \text{year}) + (\text{age}\vert \text{individual})$ | 165.72 ± 60.18 | 7.120 | 1/7 | 0.008 | 58/28 |
| MAM 4 (females): $\text{breast stripe} \sim \text{FGR} + \text{Jul.date} + (1\vert \text{year}) + (\text{age}\vert \text{individual})$ | | 12.743 | 2/6 | 0.002 | 47/21 |
| FGR | 0.278 ± 0.144 | 3.414 | 1/7 | 0.065 | |
| Jul.date | -0.013 ± 0.003 | 12.309 | 1/7 | <0.001 | |
| MAM 5 (males): $\text{yellow chroma} \sim \text{HL} + (1\vert \text{year}) + (\text{age}\vert \text{individual})$ | 0.001 ± 0.001 | 2.004 | 1/6 | 0.157 | 58/28 |
| MAM 6 (females): $\text{yellow chroma} \sim \text{HL} + \text{mass} + (1\vert \text{year}) + (\text{age}\vert \text{individual})$ | | 6.687 | 2/6 | 0.035 | 46/21 |
| HL | 0.001 ± 0.001 | 3.757 | 1/7 | 0.053 | |
| mass | 0.003 ± 0.002 | 2.821 | 1/7 | 0.093 | |
| MAM 7 (males): $\text{brightness} \sim \text{age} + \text{age}^2 + \text{mass} + \text{HL} + \text{Jul.date} + \text{FGR} + \text{T} + \text{T:Jul.date} + (1\vert \text{year}) + (\text{age}\vert \text{individual})$ | | 37.744 | 8/6 | <0.001 | 58/28 |
| age | 792.570 ± 245.120 | 18.672 | 2/12 | <0.001 | |
| age<sup>2</sup> | -4631.380 ± 1387.940 | 9.485 | 1/13 | 0.002 | |
| mass | -21340.220 ± 5590.590 | 14.130 | 1/13 | <0.001 | |
| HL | -427.940 ± 166.220 | 7.077 | 1/13 | 0.008 | |
| Jul.date | 32.220 ± 16.070 | 8.397 | 2/12 | 0.015 | |
| FGR | -1570.390 ± 910.940 | 3.067 | 1/13 | 0.080 | |
| T | 5834.870 ± 1819.610 | 13.524 | 2/12 | 0.001 | |
| T:Jul.date | -40.240 ± 13.660 | 8.311 | 1/13 | 0.004 | |
| MAM 8 (females): $\text{brightness} \sim \text{age} + (1\vert \text{year}) + (\text{age}\vert \text{individual})$ | 449.800 ± 208.000 | 4.879 | 1/6 | 0.027 | 47/21 |

### Inter-annual stability of condition-related traits and their relationship with T levels

From all condition-dependent traits, only the breast stripe area (Fig. S3A-B in SI), yellow chroma (Fig. S4A-B in SI), body mass (Fig. S8A-B in SI) and the H/L ratio in females (Fig. S6B in SI) were somewhat inter-annually repeatable in absolute values, with differences between sexes and young and adult individuals (Fig. 1; Table S2A and S2B in SI). In contrast, brightness of the yellow carotenoid ornaments (Fig. S5A-B in SI), H/L ratio in males (Fig. S7A in SI), and FGR (Fig. S6A-B in SI) had no detectable repeatability. This pattern of relatively compromised absolute repeatability was in most cases in contrast to the reasonably high-rank consistency (Fig. 1 and Table S2B in SI).

Finally, we investigated whether T levels predict FGR, body mass, and the extent of ornamental trait expression. Neither FGR nor body mass were associated with T levels (MAM2; Table 1, Table S3 in SI). Also, we found no associations between the breast stripe area and T levels in either sex (MAM3 and MAM4, Table 1, Table S3 in SI), and no relationships for the carotenoid-ornament chroma (MAM5 and MAM6 in Table 1, Table S3 in SI). However, unlike in females (MAM8, Table 1, Table S3 in SI), our results showed a strong positive relationship between T levels and yellow plumage brightness in males (slope = 5834.87, *p* = 0.001, MAM7 in Table 1, Table S3 in SI and Fig. 3).

**Figure 3.**
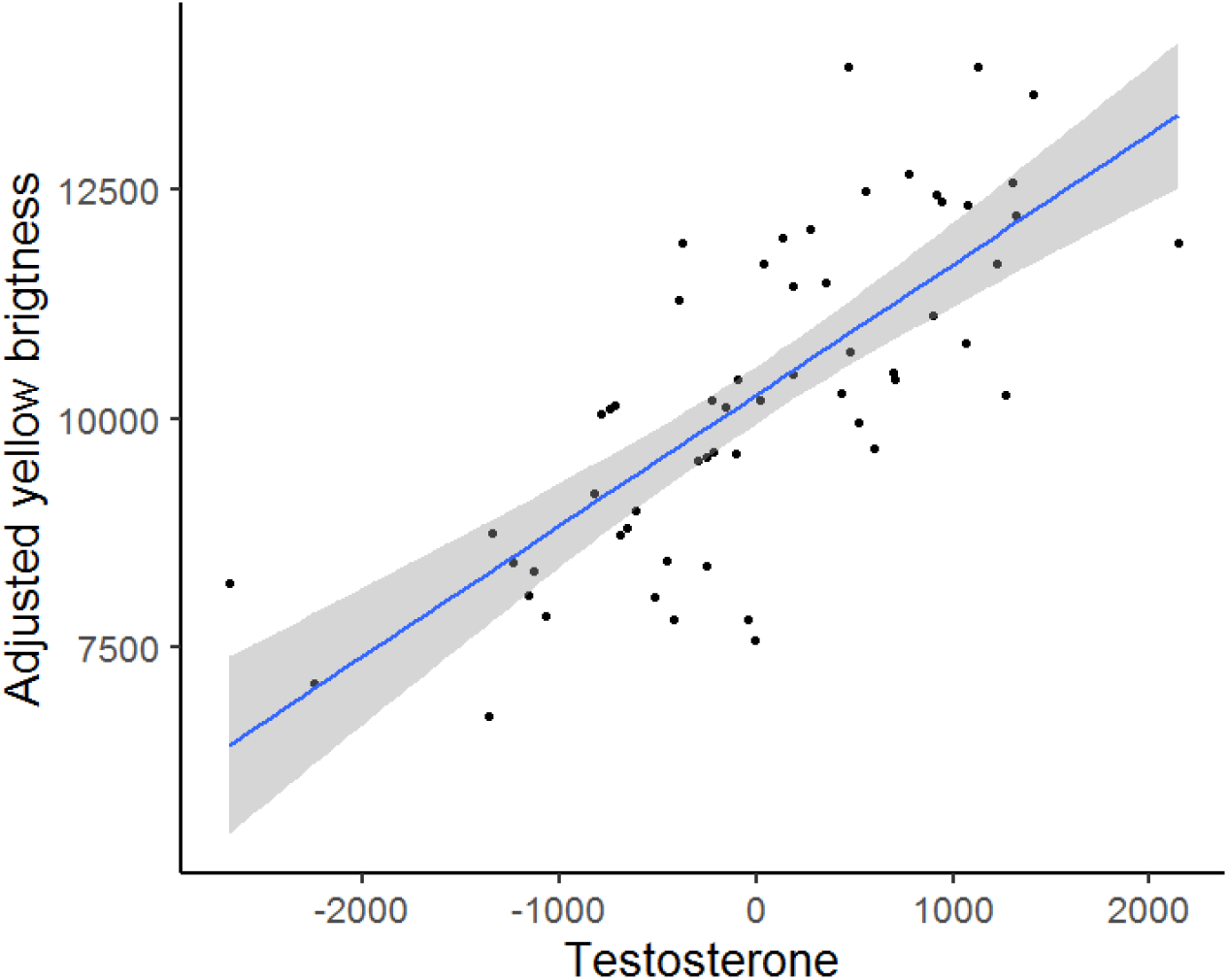
The relationship between brightness and testosterone (T) levels in repeatedly captured males (N_md_ = 28, N_obs_ = 58). The yellow brightness is shown as residuals from minimal adequate model 7 (Table 1), with the confidence interval indicated by the grey shaded area

## Discussion

In this study, we found for the first time that plasma T levels in birds are inter-annually stable in both sexes, yet with a higher degree of stability in females. Male T levels showed a quadratic dependence on age with a midlife peak. In contrast to males, female T levels were generally stable throughout their lifetime. We did not find any relationships between T levels and most condition-related traits tested, although most of these traits were also inter-annually rank-stable. The only significant and strong positive association was found between male T levels and brightness of the yellow carotenoid-based plumage ornaments, presumably serving as an honest signal in sexual selection.

Given the demanding sampling design of longitudinal studies, inter-annual plasma T level stability had not been properly tested in birds. This study provides the first evidence that avian plasma T levels are inter-annually stable in both sexes. The relatively surprising finding of higher inter-annual T stability in females compared to males appears to result from ontogenetic fluctuations in male T levels. This is supported by the finding that male T repeatability increased when the observations of young (two-year-old) individuals were excluded, indicating higher T stability after reaching reproductive maturity. However, the lower differences between sexes in individual T consistency ranks indicate that males maintained their relative ranks in the population rather than absolute plasma T levels. Similarly, T levels have been found to be inter-annually stable in the plasma of human women (Burger et al., 2000) and beluga whales (Montano, Robeck, Steinman, & O’Brien, 2017) and lesser mouse lemur (*Microcebus murinus*) males (Perret, 1992). Furthermore, also consistent with our results, higher T levels in ontogeny appear to be linked with lower T stability, as seen in a dataset of pubertal boys (Klipker, Wrzus, Rauers, Boker, & Riediger, 2017). In males, steep increases in T levels towards peak levels followed by steep decreases impair repeatability in the data, in contrast to females with relatively flat T trajectories.

In great tit males, we observed a polynomic dependence of T levels on age, with an initial three-fold increase, a peak in middle-aged individuals (minimum estimated age of four years), and a steadily decreasing trend later in life. In birds, the production of various hormones has been suggested to decline with age, co-occurring with a decrease in reproductive output (Ottinger, 1996). However, our results in males are well consistent with longitudinal data in humans (Vermeulen et al. 1972; Harman et al. 2001), as well as in long-lived Florida scrub-jays (*Aphelocoma coerulescens),* with plasma T levels peaking in middle-age (in jays there was also a ca. three-fold initial increase in T levels followed by a decrease towards the minimum T levels in the oldest, yet fertile birds; Wilcoxen et al. 2013). Although no other current avian study has adopted a longitudinal design that prevents the risk of biased interpretations due to selective disappearance in different age cohorts, further support for our results can be found in some cohort studies. In short-lived laboratory-kept Japanese quails (*Coturnix japonica;* Balthazart et al. 1984), plasma T levels dropped almost two times in the oldest age class compared to younger ones. Similarly, in longed-lived common terns (*Sterna hirundo),* male plasma T levels first sharply increased and reached a peak at maturity (2-5 years), the started to moderately decrease with age (Nisbet et al., 1999). An analogous trend suggesting the effects of hormonal senescence was also described for feather T levels in barn swallows (Adámková et al. 2019). These results are in contrast with most other avian studies (none using a longitudinal design), which have not detected any age-related changes in male T levels. The lack of such a relationship in studies comparing young vs adult individuals (e.g. in the house finch, *Haemorhous mexicanus,* Belthoff et al. 1994; the Florida scrub-jay *Aphelocoma coerulescens,* Schoech et al. 1996 and the magnificent frigatebird, *Fregata magnificens,* Madsen et al. 2007) may have been caused, for example, by the delayed breeding of younger birds at the time of their capture. Other studies with no detected relationship between age and T using cross-sectional designs (e.g. in the cliff swallow; *Petrochelidon pyrrhonota;* Smith et al. 2005), and the superb fairy-wren; *Malurus cyaneus;* Peters et al. 2002) may have been biased by selective disappearance (Zhang, Vedder, Becker, & Bouwhuis, 2015). Thus, despite previous controversies arising from cohort-based studies, our data support the universal male ontogenetic patterns in T levels across birds and mammals.

While physiologically important (Goymann & Wingfield, 2014), no longitudinal study has yet examined whether female T undergoes age-related changes through ontogeny in birds. Recently, female T levels have been shown to decline with age in two passerines, the pied flycatcher (*Ficedula hypoleuca;* Moreno et al. 2014) and the barn swallow (Adámková et al., 2019) both using cross-sectional data. In contrast, in our study female plasma T levels remained stable during their lifetime. Other longitudinal studies are needed to resolve whether there are ontogenetic parallels between mammals and birds in female T levels.

Unlike yellow plumage brightness, FGR and male H/L, the body mass and female H/L as well as the black breast stripe area and yellow plumage chroma were shown to be repeatable. This likely partially corresponds with stronger genetic components, especially in melanin-based ornaments (Grunst, Rotenberry, & Grunst, 2014; Senar & Quesada, 2006). The absence of repeatability in brightness may reflect its higher phenotypic plasticity with strong seasonal effects (Senar, Figuerola, & Pascual, 2002) but see Senar and Quesada, (2006).

Elevated T levels induce higher aggressivity, mating, and explanatory behaviours with more social interactions, all of which may interfere with body condition and increase physiological stress (Braude, Tang-Martinez, & Taylor, 1999) for the benefit of increased fitness, and reproduction-related traits including ornaments. Despite links between T levels and several condition-dependent traits being widely described in other studies (for evidence under the Immunocompetence hypothesis; Folstad and Karter 1992 or the Oxidation handicap hypothesis; Alonso-Alvarez et al. 2008, see e.g. Evans et al. 2000; Galván and Alonso-Alvarez 2010; Duckworth et al. 2004), in our study, we found only limited evidence supporting these associations. We did not observe the predicted negative link between T and body condition or the positive relationship between T levels and the H/L ratio, an indicator of physiological stress. This is, however, consistent with several other studies in passerine birds (Buchanan et al. 2003; Seddon and Klukowski 2012).

Similarly, little support was found for links between T and plumage ornaments. This may result from the fact that the effects of T on bird coloration are often complex and condition-dependent, and can therefore adopt variable directions or remain entirely lacking (Fargallo et al. 2007; Alonso-Alvarez et al. 2009; Rull et al. 2016). Furthermore, plasma T levels influence pigment deposition into ornaments only during the moulting period in the previous season (Kimball, 2006), so the absence of these relationships may also result from low individual inter-seasonal T rank stability, which remains unknown (Kempenaers et al. 2008; but see Adámková et al. 2019). However, we observed a strong positive association between T plasma levels and the brightness of the yellow plumage ornament in re-captured males. We hypothesise that this relationship may reflect the production of preen oils that are spread on feathers as protection against water and parasites (Moreno-Rueda, 2017). Uropygial gland preen oil secretion has been shown to positively correlate with plasma T levels (Amet, Abalain, Daniel, Di Stefano, & Floch, 1986; Floch, Floch, Morfin, & Daniel, 1988), which can influence the spectral structural properties of feathers (Møller & Mateos-González, 2018). Importantly, higher brightness indicates individual quality in tits (Senar et al., 2002).

Our findings showing that avian plasma T levels are inter-annually stable in both sexes provide an important basis for further research linking hormonal levels with other physiological and life-history traits. The individual rank consistency validates the use of temporarily-variable T levels to explain hormonal effects in traits measured at different time points during the lifetime, which is particularly important when investigating condition-related traits in cross-cohort datasets. Nevertheless, consistent with mammals our results also demonstrated age-dependent changes in T levels in males, indicating that this pattern may be universal across vertebrates. Since the inter-annual stability in condition-related traits mostly lacked any clear relationship to plasma T concentrations in our study, we suggest that a detailed description of the regulatory effects of T in correlative studies may require more detailed information on the individual histories of investigated individuals. This research opens the way to further understanding of context-dependency and variation in the hormonal regulation of fitness-related trait expression.

## Supporting information

Supplementary Fille 1

## Acknowledgement

We thank Hana Pinkasová, Jitka Vinklerová, Sylvie Dlugošová, Daniel Divín, Julie Vacková, Petra Blahutová, Petra Špatenková, Lenka Pelikánová, Barbora Stolínová, Monika Dvořáková, Martin Rychlý for their assistance with field sampling and Barbora Bílková, Zuzana Świderská and Tomáš Vrkoslav for their help in laboratory. We are also grateful to David Haderkopf for language corrections. This study was supported by Charles University (grants Nos. GAUK 1158217, PRIMUS/17/SCI/12 and UNCE 204069), the Czech Science Foundation (projects P502/19-20152Y and 15-11782S), Institutional Research Support No. 260571/2020) and by the Grant Agency of the Faculty of Environmental Sciences at the Czech University of Life Sciences Prague (projects IGA 20144268 and IGA 20154214). The authors declare no conflict of interest. The experiment was approved by the Environmental Protection Department of Prague City Hall (permit No. S-MHMP-1061728/2010/OPP-V-790/R-235/Bu) and by the Ethical Committee of the Institute of Vertebrate Biology, Czech Academy of Sciences (Permit No. 09/2015). Permission for bird capturing and ringing was granted by the Bird Ringing Centre of the National Museum in Prague.

## Authors’ contributions

M.T., M.V., and T.A. designed the study. M.T., T.K., H.V., J.S., H.P., P.B., J.K., T.A., and M.V. collected samples. M.T., T.K., and J.E. performed the laboratory analysis. M.T. and M.V. interpreted the data and drafted the manuscript. All authors contributed with their comments to the final approved version of the manuscript.

## Data availability statement

The data supporting the results are archived in the Dryad repository: https://datadryad.org/stash/share/dqno1iviZi4v00C3nfBWlMwZUWkVsl3rzCaYrn3JgCc

## References

Adámková, M., Bílková, Z., Tomášek, O., Šimek, Z., & Albrecht, T. (2019). Feather steroid hormone concentrations in relation to age, sex, and molting time in a long-distance migratory passerine. Ecology and Evolution, 9(16), 9018–9026. doi: 10.1002/ece3.5447

Albrecht, T., Vinkler, M., Schnitzer, J., Polakova, R., Munclinger, P., & Bryja, J. (2009). Extra-pair fertilizations contribute to selection on secondary male ornamentation in a socially monogamous passerine. Journal of Evolutionary Biology, 22(10). 2020–2030. doi: 10.1111/j.1420-9101.2009.01815.x

Alonso-Alvarez, C., Pérez-Rodríguez, L., Garcia, J. T., & Viñuela, J. (2009). Testosterone-mediated trade-offs in the old age: A new approach to the immunocompetence handicap and carotenoid-based sexual signalling. Proceedings of the Royal Society B: Biological Sciences, 276(1664), 2093–2101. doi: 10.1098/rspb.2008.1891

Amet, Y., Abalain, J. H., Daniel, J. Y., Di Stefano, S., & Floch, H. H. (1986). Testosterone regulation of androgen receptor levels in the uropygial gland of quails (Coturnix coturnix): A further proof for the androgen dependency of the uropygial gland. General and Comparative Endocrinology, 62(2), 210–216. doi: 10.1016/0016-6480(86)90111-5

Balthazart, J., Turek, R., & Ottinger, M. A. (1984). Altered brain metabolism of testosterone is correlated with reproductive decline in aging quail. Hormones and Behavior, 18(3), 330–345. doi: 10.1016/0018-506X(84)90020-5

Bates, D., Mächler, M., Bolker, Benjamin, M., & Walker, Steven, C. (2015). Fitting Linear Mixed-Effects Models Using lme4. Journal of Statistical Software, 67(1). doi: 10.18637/jss.v067.i01

Bauerová, P., Krajzingrová, T., Těšický, M., Velová, H., Hraníček, J., Musil, S., … Vinkler, M. (2020). Longitudinally monitored lifetime changes in blood heavy metal concentrations and their health effects in urban birds. Science of the Total Environment, 723. doi: 10.1016/j.scitotenv.2020.138002

Bauerová, P., Vinklerová, J., Hraníček, J., Čorba, V., Vojtek, L., Svobodová, J., & Vinkler, M. (2017). Associations of urban environmental pollution with health-related physiological traits in a free-living bird species. Science of the Total Environment, 601-602, 1556–1565. doi: 10.1016/j.scitotenv.2017.05.276

Belthoff, J. R., Dufty, A. M., & Gauthreaux, S. A. (1994). Plumage Variation, Plasma Steroids and Social Dominance in Male House Finches. The Condor, 96(3), 614–625. doi: 10.2307/1369464

Bernstein, R. M., Setchell, J. M., Verrier, D., & Knapp, L. A. (2012). Maternal Effects and the Endocrine Regulation of Mandrill Growth. American Journal of Primatology, 74(10), 890–900. doi: 10.1002/ajp.22038

Bouwhuis, S., Sheldon, B. C., Verhulst, S., & Charmantier, A. (2009). Great tits growing old: Selective disappearance and the partitioning of senescence to stages within the breeding cycle. Proceedings of the Royal Society B: Biological Sciences, 276(1668), 2769–2777. doi: 10.1098/rspb.2009.0457

Braude, S., Tang-Martinez, Z., & Taylor, G. T. (1999). Stress, testosterone, and the immunoredistribution hypothesis. Behavioral Ecology, 10(3), 345–350. doi: 10.1093/beheco/10.3.345

Buchanan, K. L., Evans, M. R., & Goldsmith, A. R. (2003). Testosterone, dominance signalling and immunosuppression in the house sparrow, Passer domesticus. Behavioral Ecology and Sociobiology, 55(1), 50–59. doi: 10.1007/s00265-003-0682-4

Burger, H. G., Dudley, E. C., Cui, J., Dennerstein, L., Hopper, J. L., & Henry, P. (2000). A Prospective Longitudinal Study of Serum Testosterone, Dehydroepiandrosterone Sulfate, and Sex Hormone-Binding Globulin Levels through the Menopause Transition. Journal of Clinical Endocrinology and Metabolism, 85(8), 2832–2838. doi: 10.1210/jcem.85.8.6740

Chahal, H. S., & Drake, W. M. (2007). The endocrine system and ageing. Journal of Pathology, 211(2), 173–180. doi: 10.1002/path

Crawley, M. J. (2013). The R book. Retrieved from http://www.books24x7.com/marc.asp?bookid=51275

Davis, A. K., Maney, D. L., & Maerz, J. C. (2008). The use of leukocyte profiles to measure stress in vertebrates: A review for ecologists. Functional Ecology, 22(5), 760–772. doi: 10.1111/j.1365-2435.2008.01467.x

Deviche, P., Beouche-Helias, B., Davies, S., Gao, S., Lane, S., & Valle, S. (2014). Regulation of plasma testosterone, corticosterone, and metabolites in response to stress, reproductive stage, and social challenges in a desert male songbird. General and Comparative Endocrinology, 203, 120–131. doi: 10.1016/j.ygcen.2014.01.010

Duckworth, R. A., Mendonça, M. T., & Hill, G. E. (2004). Condition-dependent sexual traits and social dominance in the house finch. Behavioral Ecology, 15(5), 779–784. doi: 10.1093/beheco/arh079

Edler, A. U., & Friedl, T. W. P. (2010). Individual quality and carotenoid-based plumage ornaments in male red bishops (Euplectes orix): Plumage is not all that counts. Biological Journal of the Linnean Society, 99(2), 384–397. doi: 10.1111/j.1095-8312.2009.01354.x

Evans, M. R., Goldsmith, A. R., & Norris, S. R. A. (2000). The effects of testosterone on antibody production and plumage coloration in male house sparrows (Passer domesticus). Behavioral Ecology and Sociobiology, 47(3), 156–163. doi: 10.1007/s002650050006

Fargallo, J. A., Martínez-Padilla, J., Toledano-Díaz, A., Santiago-Moreno, J., & Dávila, J. A. (2007). Sex and testosterone effects on growth, immunity and melanin coloration of nestling Eurasian kestrels. Journal of Animal Ecology, 76(1), 201–209. doi: 10.1111/j.1365-2656.2006.01193.x

Floch, J. Y., Floch, H. H., Morfin, R. F., & Daniel, J. Y. (1988). Testosterone Metabolism and Its Testosterone-Dependent Activation in the Uropygial Gland of Quail. Endocrine Research, 14(1), 93–107. doi: 10.1080/07435808809036342

Foerster, K., Poesel, A., Kunc, H., & Kempenaers, B. (2002). The natural plasma testosterone profile of male blue tits during the breeding season and its relation to song output. Journal of Avian Biology, 33(3), 269–275. doi: 10.1034/j.1600-048X.2002.330309.x

Folstad, I., & Karter, A. J. (1992). Parasites, bright males, and the immunocompetence handicap. American Naturalist, 139(3), 603–622. doi: 10.1086/285346

Fusani, L. (2008). Testosterone control of male courtship in birds. Hormones and Behavior, 54(2), 227–233. doi: 10.1016/j.yhbeh.2008.04.004

Galván, I., & Alonso-Alvarez, C. (2010). Yolk testosterone shapes the expression of a melanin-based signal in great tits: An antioxidant-mediated mechanism? Journal of Experimental Biology, 213(18), 3127–3130. doi: 10.1242/jeb.045096

Gonzalez, G., Sorci, G., Smith, L. C., & Lope, F. (2001). Testosterone and sexual signalling in male house sparrows (Passer domesticus). Behavioral Ecology and Sociobiology, 50(6), 557–562. doi: 10.1007/s002650100399

Goymann, W., & Wingfield, J. C. (2014). Male-to-female testosterone ratios, dimorphism, and life history - What does it really tell us? Behavioral Ecology, 25(4), 685–699. doi: 10.1093/beheco/aru019

Griffith, S. C., Parker, T. H., & Olson, V. A. (2006). Melanin-versus carotenoid-based sexual signals: is the difference really so black and red? Animal Behaviour, 71, 749–763. doi: 10.1016/j.anbehav.2005.07.016

Grubb, C. T. (2006). Ptilochronology: Feather Time and the Biology of Birds (Vol. 15). Oxford University Press.

Grunst, A. S., Rotenberry, J. T., & Grunst, M. L. (2014). Age-dependent relationships between multiple sexual pigments and condition in males and females. Behavioral Ecology, 25(2), 276–287. doi: 10.1093/beheco/art124

Guindre-Parker, S., & Love, O. P. (2014). Revisiting the condition-dependence of melanin-based plumage. Journal of Avian Biology, 45(1), 29–33. doi: 10.1111/j.1600-048X.2013.00190.x

Harman, S. M., Metter, E. J., Tobin, J. D., Pearson, J., & Blackman, M. R. (2001). Longitudinal effects of aging on serum total and free testosterone levels in healthy men. Journal of Clinical Endocrinology and Metabolism, 86(2), 724–731. doi: 10.1210/jcem.86.2.7219

Hau, M., & Goymann, W. (2015). Endocrine mechanisms, behavioral phenotypes and plasticity: known relationships and open questions From New Perspectives in Behavioural Development: Adaptive Shaping of Behaviour over a Lifetime? Frontiers in Zoology, 12(Suppl 1), 1–15. doi: 10.1186/1742-9994-12-S1-S7

Hegyi, G., Szigeti, B., Torok, J., & Eens, M. (2007). Melanin, carotenoid and structural plumage ornaments: information content and role in great tits Parus major. Journal of Avian Biology, 38(6), 698–708. doi: 10.1111/j.2007.0908-8857.04075.x

Hernández-López, L., Cerda-Molina, A. L., Díaz-Díaz, G., Chavira-Bolaños, R., & Mondragón-Ceballos, R. (2012). Aging-related reproductive decline in the male spider monkey (Ateles geoffroyi). Journal of Medical Primatology, 41(2), 115–121. doi: 10.1111/j.1600-0684.2011.00528.x

Hirschenhauser, K., Mostl, E., & Kotrschal, K. (1999). Within-pair testosterone covariation and reproductive output in Greylag Geese Anser anser. Ibis, 141(4), 577–586. doi: 10.1111/j.1474-919x.1999.tb07365.x

Kempenaers, B., Peters, A., & Foerster, K. (2008). Sources of individual variation in plasma testosterone levels. Philosophical Transactions of the Royal Society B: Biological Sciences, 363(1497), 1711–1723. doi: 10.1098/rstb.2007.0001

Ketterson, E. D., Nolan, V., & Sandell, M. (2005). Testosterone in females: Mediator of adaptive traits, constraint on sexual dimorphism, or both? American Naturalist, 166(4 SUPPL.), S85–98. doi: 10.1086/444602

Kimball, R. T. (2006). Hormonal Control of Coloration. in Bird Coloration I: measurements and mechanism. (G. E. Hill and K. McGraw, Ed.). Cambridge: Harvard University Press.

Klipker, K., Wrzus, C., Rauers, A., Boker, S. M., & Riediger, M. (2017). Within-person changes in salivary testosterone and physical characteristics of puberty predict boys’ daily affect. Hormones and Behavior, 95(August), 22–32. doi: 10.1016/j.yhbeh.2017.07.012

Madsen, V., Valkiunas, G., Iezhova, T. A., Mercade, C., Sanchez, M., & Osorno, J. L. (2007). Testosterone levels and gular pouch coloration in courting magnificent frigatebird (Fregata magnificens): Variation with age-class, visited status and blood parasite infection. Hormones and Behavior, 51(1), 156–163. doi: 10.1016/j.yhbeh.2006.09.010

Møller, A. P., & Mateos-González, F. (2018). Plumage brightness and uropygial gland secretions in barn swallows. Current Zoology, 65(2), 177–182. doi: 10.1093/cz/zoy042

Montano, G. A., Robeck, T. R., Steinman, K. J., & O’Brien, J. K. (2017). Circulating anti-Müllerian hormone concentrations in relation to age and season in male and female beluga (Delphinapterus leucas). Reproduction, Fertility and Development, 29(8), 1642–1652. doi: 10.1071/RD15537

Montgomerie, R. (2006). Analyzing colors. In: Bird Coloration I: Mechanisms and Measurements. (G. E. Hill & K. J. McGraw, Eds.). Cambridge: Harvard University Press.

Moreno-Rueda, G. (2017). Preen oil and bird fitness: a critical review of the evidence. Biological Reviews, 92(4), 2131–2143. doi: 10.1111/brv.12324

Moreno, J., Gil, D., Cantarero, A., & López-Arrabé, J. (2014). Extent of a white plumage patch covaries with testosterone levels in female Pied Flycatchers Ficedula hypoleuca. Journal of Ornithology, 155(3), 639–648. doi: 10.1007/s10336-014-1046-8

Nisbet, I. C. T., Finch, C. E., Thompson, N., Russek-Cohen, E., Proudman, J. A., & Ottinger, M. A. (1999). Endocrine patterns during aging in the common tern (Sterna hirundo). General and Comparative Endocrinology, 114(2), 279–286. doi: 10.1006/gcen.1999.7255

Ottinger, M. A. (1996). Aging in the avian brain: Neuroendocrine considerations. Seminars in Avian and Exotic Pet Medicine, 5(3), 172–177. doi: 10.1016/s1055-937x(96)80006-5

Perret, M. (1992). Environmental and social determinants of sexual function in the male lesser mouse lemur (Microcebus murinus). Folia Primatologica; International Journal of Primatology, 59(1), 1–25. doi: 10.1159/000156637

Peters, A., Delhey, K., Goymann, W., & Kempenaers, B. (2006). Age-dependent association between testosterone and crown UV coloration in male blue tits (Parus caeruleus). Behavioral Ecology and Sociobiology, 59(5), 666–673. doi: 10.1007/s00265-005-0095-7

Peters, Anne, Cockburn, A., & Cunningham, R. (2002). Testosterone treatment suppresses paternal care in superb fairy-wrens, Malurus cyaneus, despite their concurrent investment in courtship. Behavioral Ecology and Sociobiology, 51(6), 538–547. doi: 10.1007/s00265-002-0472-4

Pinheiro, J., Bates, D., DebRoy, S., & Sarkar, D. (2019). {nlme}: Linear and Nonlinear Mixed Effects Models. Retrieved from https://cran.r-project.org/package=nlme

Quesada, J., & Senar, J. C. (2006). Comparing plumage colour measurements obtained directly from live birds and from collected feathers: The case of the great tit Parus major. Journal of Avian Biology, 37(6), 609–616. doi: 10.1111/j.0908-8857.2006.03636.x

Rull, I. L., Nicolás, L., Neri-Vera, N., Argáez, V., Martínez, M., & Torres, R. (2016). Assortative mating by multiple skin color traits in a seabird with cryptic sexual dichromatism. Journal of Ornithology, 157(4), 1049–1062. doi: 10.1007/s10336-016-1352-4

Schindelin, J., Rueden, C. T., Hiner, M. C., & Eliceiri, K. W. (2015). The ImageJ ecosystem: An open platform for biomedical image analysis. Molecular Reproduction and Development, 82(7–8), 518–529. doi: 10.1002/mrd.22489

Schoech, S. J., Mumme, R. L., & Wingfield, J. C. (1996). Delayed breeding in the cooperatively breeding Florida scrubjay (Aphelocoma coerulescens): Inhibition or the absence of stimulation? Behavioral Ecology and Sociobiology, 39(2), 77–90. doi: 10.1007/s002650050269

Seddon, R. J., & Klukowski, M. (2012). Influence of Stressor Duration on Leukocyte and Hormonal Responses in Male Southeastern Five-Lined Skinks (Plestiodon inexpectatus). Journal of Experimental Zoology Part A: Ecological Genetics and Physiology, 317(8), 499–510. doi: 10.1002/jez.1742

Senar, J. C., Figuerola, J., & Pascual, J. (2002). Brighter yellow blue tits make better parents. Proceedings of the Royal Society B: Biological Sciences, 269(1488), 257–261. doi: 10.1098/rspb.2001.1882

Senar, J. C., & Quesada, J. (2006). Absolute and relative signals: A comparison between melanin-and carotenoid-based patches. Behaviour, 143(5), 589–595. doi: 10.1163/156853906776759484

Smith, L. C., Raouf, S. A., Bomberger Brown, M., Wingfield, J. C., & Brown, C. R. (2005). Testosterone and group size in cliff swallows: Testing the “challenge hypothesis” in a colonial bird. Hormones and Behavior, 47(1), 76–82. doi: 10.1016/j.yhbeh.2004.08.012

Stoffel, M. A., Nakagawa, S., & Schielzeth, H. (2017). rptR: repeatability estimation and variance decomposition by generalized linear mixed-effects models. Methods in Ecology and Evolution, 8(11), 1639–1644. doi: 10.1111/2041-210X.12797

Svensson, L., & Baker, K. (1992). Identification guide to European passerines (4th ed.). Stockholm: British Trust for Ornithology.

Svobodová, J., Bauerová, P., Eliáš, J., Velová, H., Vinkler, M., & Albrecht, T. (2018). Sperm variation in Great Tit males (Parus major) is linked to a haematological health-related trait, but not ornamentation. Journal of Ornithology, 159(3), 815–822. doi: 10.1007/s10336-018-1559-7

Svobodová, J., Gabrielová, B., Synek, P., Marsik, P., Vaněk, T., Albrecht, T., & Vinkler, M. (2013). The health signalling of ornamental traits in the Grey Patridge (Perdix perdix). Journal of Ornithology, 154(3), 717–725. doi: 10.1007/s10336-013-0936-5

Van Duyse, E., Pinxten, R., & Eens, M. (2003). Seasonal fluctuations in plasma testosterone levels and diurnal song activity in free-living male great tits. General and Comparative Endocrinology, 134(1), 1–9. doi: 10.1016/S0016-6480(03)00213-2

Vermeulen, A., Rubens, R., & Verdonck, L. (1972). Testosterone secretion and metabolism in male senescence. Journal of Clinical Endocrinology and Metabolism, 34(4), 730–735. doi: 10.1210/jcem-34-4-730

Vinkler, M., Schnitzer, J., Munclinger, P., & Albrecht, T. (2012). Phytohaemagglutinin skin-swelling test in scarlet rosefinch males: low-quality birds respond more strongly. Animal Behaviour, 83(1), 17–23. doi: 10.1016/j.anbehav.2011.10.001

Weaver, R. J., Santos, E. S. A., Tucker, A. M., Wilson, A. E., & Hill, G. E. (2018). Carotenoid metabolism strengthens the link between feather coloration and individual quality. Nature Communications, 9(1). doi: 10.1038/s41467-017-02649-z

Wilcoxen, T. E., Bridge, E. S., Boughton, R. K., Hahn, T. P., & Schoech, S. J. (2013). Physiology of reproductive senescence in Florida scrub-jays: Results from a long-term study and GnRH challenge. General and Comparative Endocrinology, 194, 168–174. doi: 10.1016/j.ygcen.2013.09.016

Wolf, T. E., Schaebs, F. S., Bennett, N. C., Burroughs, R., & Ganswindt, A. (2018). Age and socially related changes in fecal androgen metabolite concentrations in free-ranging male giraffes. General and Comparative Endocrinology, 255, 19–25. doi: 10.1016/j.ygcen.2017.09.028

Zhang, H., Vedder, O., Becker, P. H., & Bouwhuis, S. (2015). Age-dependent trait variation: The relative contribution of within-individual change, selective appearance and disappearance in a long-lived seabird. Journal of Animal Ecology, 84(3), 797–807. doi: 10.1111/1365-2656.12321

