## Supplementary Fille 1 for "Inter-annual stability and age-dependent changes in plasma testosterone levels in a longitudinally monitored free-living passerine bird"

**Supporting Information**

**Supplementary materials and methods online (SMMO)**

**SMMO 1: Field procedures**

Between 2012 and 2017 in total 49 individuals of the great tit (28 males and 21 females) were repeatedly sampled in at least two or three subsequent breeding seasons (April–May; for age-structure see; see Fig. S1). The study was conducted in a deciduous city forest at the edge of Prague (Czech Republic; 50°8′10.591″N, 14°27′51.144″E, ~ 315-360 m above the sea level, total area ca. 0.9 km$^2$) on a free-living population breeding in nest boxes (for a detailed description of the study locality and field procedures, see Svobodová et al. 2018 and Bauerová et al. 2020). Briefly, the study area is located in the vicinity of a meteorological station Kbely (Czech Hydrometeorological Institute) from where information on mean air temperature seven days before catching (further referred to as temperature) was obtained. All birds were captured when the nestlings were 7-14 days old by placing a mist net against the nest box opening (following the standard protocol of the Czech Bird Ringing Centre). Immediately after capture, approximately 150 μl of blood was collected by an insulin syringe (Omican 50-50I.U./0,5ML 30G x12; B. Braun, Melsungen, Germany) from a jugular vein of each individual. A blood smear was prepared from a drop of blood, part of the sample was stored frozen in a microtube with 96% ethanol for later use (as genetic material and the material for heavy metal analysis; not used in this study) and a subsample of the blood (ca. 70-100 μl) was stored unfrozen in a cool box and later the same day transported into the laboratory. Here each blood sample was centrifuged in a microcentrifuge (type 5424, Eppendorf, Hamburg, Germany) at 8000 rpm for 5 minutes and the obtained plasma was frozen at −80 °C. In the field, weight (measured by a digital scale, $d$ = 0.02 g, type PPS200, Pesola, Schindellegi. Switzerland) and tarsus length (measured by a digital calliper, accuracy 0.01 mm; Kinex, Prague, Czech Republic) were recorded. We later calculated size-standardised body mass as the ratio of weight to tarsus length (further referred to as body mass). The second tail rectrix from the left side was collected from each individual for the ptilochronological assessment of FGR (Grubb, 2006). To assess melanin-based and yellow carotenoid-based plumage ornamentation, we first, collected a standardised digital image of the black breast stripe using a scanner (type Perfection V30, Seiko Epson, Nagano, Japan). The procedure was performed in a mobile dark tent, with grey and colour standard reference swatches equipped with a ruler (GC 18 grey card and Q 14 colour and grey chart; Danes-Picta, Prague, Czech Republic) in a standardised position of the bird. Second, samples of ornamental feathers from the upper part of the yellow breast area were collected for light spectral analysis (ca. 20-25 feathers). Finally, each individual was tagged with a steel ring with a unique code of the Czech Bird Ringing Centre, National Museum in Prague. In this long-term study, the minimum age of each bird was estimated based on a combination of the ringing records and plumage age characteristics at their first capture (the colour differences in primary and secondary covers allows distinguishing between the second year and older individuals; Svensson and Baker 1992).

All applicable international, national, and/or institutional guidelines for the care and use of animals were followed. The experiment was approved by the Environmental Protection Department of Prague City Hall (permit No. S-MHMP-1061728/2010/OPP-V-790/R-235/Bu) and by the Ethical Committee of the Institute of Vertebrate Biology, Czech Academy of Sciences (Permit No. 09/2015). Permission for catching and ringing adult birds was granted by the Bird Ringing Centre of the National Museum in Prague.

**SMMO 2: Plasma testosterone assay**

Plasma T was quantified using Testosterone ELISA Kit (480 solid well; product No 582701; Cayman Chemical Company, Ann Arbor, USA) according to the manufacturer's instructions with measuring range 3.9-500 pg/ml and sensitivity 6 pg/ml for T. As an input, 20 μl of plasma sample was diluted 5.5×. The samples were assayed in duplicates with samples from a single individual assayed on the same plate. Following manufacturer's instructions, on each plate T standards allow inferring T concentration from the calibration curve. We calculated the measurement repeatability from replicates, intra-assay variation as the percent variation coefficient (CV) between replicates, and inter-assay variation as the percent CV between one pooled sample assayed on all the plates. The repeatability of measurement between duplicates was $r = 0.91$ ($p < 0.001$) and intra- and inter-assay (CV) were 13.01 (N =102) and 24.01 % (N = 6), respectively.

**SMMO 3: Haematological analysis**

We adopted the haematological procedures previously reported by Vinkler et al. (2010). Briefly, from each blood sample collected into a heparinised insulin syringe, a blood smear was prepared from a small droplet of blood in the field. Dried blood smears were later stained with the Wright-Giemsa Modified stain (product No. WG128, Sigma-Aldrich, St. Luis, MO, USA) and used to assess differential leukocyte counts under a light microscope with 100× objective magnification (Olympus Corporation, Tokyo, Japan, type CX-31). For the differential leukocyte count, 120-130 leukocytes per individual were counted, differentiating five cell types: lymphocytes, monocytes, heterophils, basophils, and eosinophils. Out of these data, we further calculated the H/L ratio. All cell counting was performed by one person only (TK) to minimise any potential variability among the measurements.

**SMMO 4: Ptilochronological analysis**

To evaluate the nutritional status of the birds in the time of moulting, we performed a ptilochronological analysis of FGR using the second left rectrix (Vinkler, Schnitzer, Munclinger, & Albrecht, 2012). First, we scanned all second rectrices with a 50 mm scale using a scanner (Epson V30) in the greyscale reflex mode with 600dpi resolution. Then we processed the images in Corel Photo-Paint X3 software (Corel Corporation, Ottawa, Canada) using the function of Local Equalization (parameters: Width 100 and Height 100). Subsequently, we measured the total rectrix length and estimated the FGR as the mean growth bar width in a segment of 10 growth bars with the centre located at 2/3 of the feather length using the ImageJ software (v. 2.0.0; The National Institutes of Health, Bethesda, Maryland, USA, Schindelin et al. 2015).

**SMMO 5: Analysis of the plumage ornament**

The digital images of the great tit black breast stripe area were analysed using Adobe PHOTOSHOP CS. 2 software (v. 10.0; Adobe Systems, San Jose, USA) according to Bauerová et al. (2017). The total area of the black breast stripe was measured at 50 mm in length from the neck using the selection tool and area measuring function. For 20 randomly selected individuals, we checked the repeatability of the measurement that was found to be high ($r = 0.9$, $p < 0.001$).

Yellow carotenoid-based breast ornament was measured using the Avaspec 2048 spectrometer with an Avalight XE light source and the Avasoft 7.0 processing system (Avantes, Eerbeek, Netherlands). For each individual, we measured the colour of a layer of 20-25 sampled carotenoid-based feathers fixed on the surface of a glass slide with a tape (according to Quesada and Senar 2006) under standardized conditions (for more details about the method, see Svobodová et al. 2018 and Albrecht et al. 2009). For determination of yellow saturation of the yellow colouration, we calculated the chroma (the difference between reflectance at 700 nm and reflectance at 450 nm, relative to reflectance at 700nm; the interval for absorbance of carotenoid pigments is 450–700 nm) and total brightness (total reflectance, i.e. the sum of reflectance in visible light from 300 to 700 nm) from the spectral data (Montgomerie, 2006). For 20 randomly selected individuals, we checked the repeatability of the measurement that was again shown high ($r = 0.85$, $p < 0.001$).

### Tables and Figures

**Table S1** Summary statistics of raw plasma testosterone concentrations [pg/ml] in different age classes in re-captured individuals of the free-living great tits. Sex – males (M), females (F), Age – minimal estimated age based on bird ringing and plumage characteristics, $N_{obs}$ – number of observations, $N_{ind}$ – number of individuals (in total, Nobs = 105, $N_{ind}$ = 49)

| Sex | Age | N | Mean ± SD | Range |
|---|---|---|---|---|
| M | 2 | 12 | 125.50 ± 71.14 | 61.95 ± 312.45 |
| F | | 13 | 127.00 ± 85.33 | 27.50 ± 276.03 |
| M | 3 | 21 | 393.66 ± 382.59 | 35.71 ± 1570.98 |
| F | | 18 | 100.82 ± 72.80 | 25.80 ± 276.37 |
| M | 4 | 17 | 535.03 ± 446.41 | 61.62 ± 1803.47 |
| F | | 9 | 115.28 ± 85.51 | 38.62 ± 302.88 |
| M | 5 | 7 | 530.26 ± 422.45 | 131.98 ± 1296.56 |
| F | | 4 | 80.18 ± 48.18 | 41.39 ± 150.61 |
| M | 6 | 1 | 267.35 | - |
| F | | 2 | 120.96 ± 69.54 | 71.79 ± 170.13 |
| M | 7 | 0 | - | - |
| F | | 1 | 124.57 | - |

**Table S2** Inter-annual within individual repeatability (r) of testosterone (T) and condition-related traits based on (A) their absolute trait values (i.e. absolute repeatability) and (B) weighted relative ranks (i.e. individual consistency ranks) in repeatedly captured individuals of the great tit. For the individual consistency rank, the population rank is weighted by the number of observations in a given year. The calculations were done separately for both sexes with all observations and with excluded observations of young birds (two-year-old; 2Y). Testosterone – residuals for plasma testosterone, breast stripe – the black breast stripe area, yellow chroma – yellow chroma of carotenoid ornament, brightness – brightness of yellow carotenoid ornament, FGR – feather growth rate, i.e. daily increase of growth bar width of rectrices, HL – H/L ratio (number of heterophils/ number of lymphocytes), mass – body mass as the ratio of weight and tarsus length, $N_{obs}$ – number of observations, $N_{ind}$ – number of individuals. All M – all male individuals ($N_{ind}$ = 28, $N_{obs}$ = 58), +2Y M – only male individuals older than two-year-old ($N_{ind}$ = 17, $N_{obs}$ = 36), all F – all female individuals ($N_{ind}$ = 21, $N_{obs}$ = 47), +2Y F – only female individuals older than two-year-old ($N_{ind}$ = 11, $N_{obs}$ = 24). Statically significant repeatabilities at *p*-value < 0.05 are in bold

(A)

| | all M | | +2Y M | | all F | | +2Y F | |
|---|---|---|---|---|---|---|---|---|
| Trait | *r* | *p*-value | *r* | *p*-value | *r* | *p*-value | *r* | *p*-value |
| Testosterone | 0.103 | 0.304 | **0.385** | **0.045** | **0.576** | **0.001** | **0.590** | **0.016** |
| Breast stripe area | 0.058 | 0.400 | 0.110 | 0.346 | 0.174 | 0.209 | **0.481** | **0.029** |
| Yellow chroma | 0.248 | 0.110 | 0.404 | 0.053 | 0.122 | 0.312 | 0.014 | 1.000 |
| Yellow brightness | 0.000 | 1.000 | 0.000 | 1.000 | 0.000 | 1.000 | 0.000 | 1.000 |
| FGR | 0.000 | 1.000 | 0.000 | 1.000 | 0.000 | 0.500 | 0.000 | 1.000 |
| HL | 0.000 | 0.500 | 0.000 | 1.000 | 0.279 | 0.072 | 0.298 | 0.120 |
| Body mass | **0.441** | **0.006** | 0.355 | 0.061 | 0.277 | 0.090 | 0.297 | 0.147 |

(B)

| | all M | | +2Y M | | all F | | +2Y F | |
|---|---|---|---|---|---|---|---|---|
| Trait | *r* | *p*-value | *r* | *p*-value | *r* | *p*-value | *r* | *p*-value |
| Testosterone | **0.411** | **0.008** | 0.312 | 0.083 | **0.489** | **0.005** | 0.407 | 0.087 |
| Breast stripe area | 0.082 | 0.359 | 0.089 | 0.391 | **0.423** | **0.009** | **0.554** | **0.014** |
| Yellow chroma | 0.192 | 0.179 | 0.385 | 0.064 | 0.306 | 0.078 | 0.453 | 0.077 |
| Yellow brightness | 0.000 | 1.000 | 0.000 | 1.000 | 0.000 | 1.000 | 0.000 | 1.000 |
| FGR | 0.000 | 0.500 | 0.000 | 1.000 | 0.091 | 0.367 | 0.289 | 0.200 |
| HL | 0.178 | 0.200 | 0.000 | 1.000 | **0.484** | **0.005** | **0.514** | **0.028** |
| Body mass | **0.460** | **0.005** | 0.356 | 0.059 | **0.396** | **0.020** | **0.705** | **0.001** |

**Table S3** Full Linear Model (LM) and Linear Mixed Models (LMMs) tested in the great tit study. For LMMs models (M2 – M8), a year of capture and individual ID were used as random effects with random intercepts. The age was given as a random slope effect within the individual ID (age|ID). Residuals for testosterone (T) from full LM model 1 were used in all other models. $T_{raw}$ – plasma testosterone concentration, T_plate – testosterone plate (batch in which the sample was measured), handling time - the time between capturing and blood sampling, time - the time of capture, T – plasma testosterone levels as residuals from model 1, breast stripe – the black breast stripe area, individual – individual identification based on bird ringing, Sex – sex, age – minimal estimated age based on bird ringing (age^2 – polynomic term), T_ave7 – mean daily temperature 7 days before catching, FGR – feather growth rate, i.e. daily increase of growth bar width of rectrices [mm], Jul.date – Julian date from the beginning of the year, mass – body mass as the ratio of weight and tarsus length, HL – H/L ratio (number of heterophils/ number of lymphocytes), brightness – brightness of yellow carotenoid ornament, yellow chroma – yellow chroma of carotenoid ornament, $N_{obs}$ – number of observations, $N_{ind}$ – number of individuals

| **Full model** | **F** | **Df** | ***p*** | **$N_{obs}$/$N_{ind}$** |
|---|---|---|---|---|
| model 1: **log ($T_{raw}$)** ~ **T_plate** + handling time + time | 2.365 | 7/104 | 0.028 | 105/49 |
| model 2: **T** ~ sex + age + age^2 + Jul.date + tars + mass + FGR + HL + T_ave7 + sex: age^2+ sex:age +HL:sex + (1\|year) + (age\|individual) | 49.970 | 12/6 | <0.001 | 105/49 |
| model 3 (males): log (breast stripe) ~ age + age^2 + tars + mass + FGR + HL + Jul.date + **T + T:Jul.date** + (1\|year) + (age\|individual) | 12.255 | 8/7 | 0.140 | 58/28 |
| model 4 (females): breast stripe ~ age + age^2 + tars + mass + FGR + HL + Jul.date + **T + T:Jul.date** + (1\|year) + (age\|individual) | 17.194 | 9/6 | 0.046 | 47/21 |
| model 5 (males): yellow chroma ~ age + age^2 + tars + mass + FGR + HL + Jul.date + **T + T:Jul.date** + (1\|year) + (age\|individual) | 5.624 | 8/6 | 0.777 | 58/28 |
| model 6 (females): yellow chroma ~ age + age^2 + tars + mass + FGR + HL + Jul.date + **T + T:Jul.date** + (1\|year) + (age\|individual) | 12.980 | 9/6 | 0.164 | 46/21 |
| model 7 (males): brightness ~ age + age^2 + tars + mass + FGR + HL + Jul.date + **T + T:Jul.date** + (1\|year) + (age\|individual) | 37.744 | 9/6 | <0.001 | 58/28 |
| model 8 (females): brigtness ~ age + age^2 + tars + mass + FGR + HL + Jul.date + **T + T:Jul.date** + (1\|year) + (age\|individual) | 16.093 | 9/6 | 0.065 | 47/21 |

**Figure S1** A Histogram of number of observations in each age category for both sexes in the dataset of re-captured great tits (Nobs = 105, $N_{ind}$ = 49)

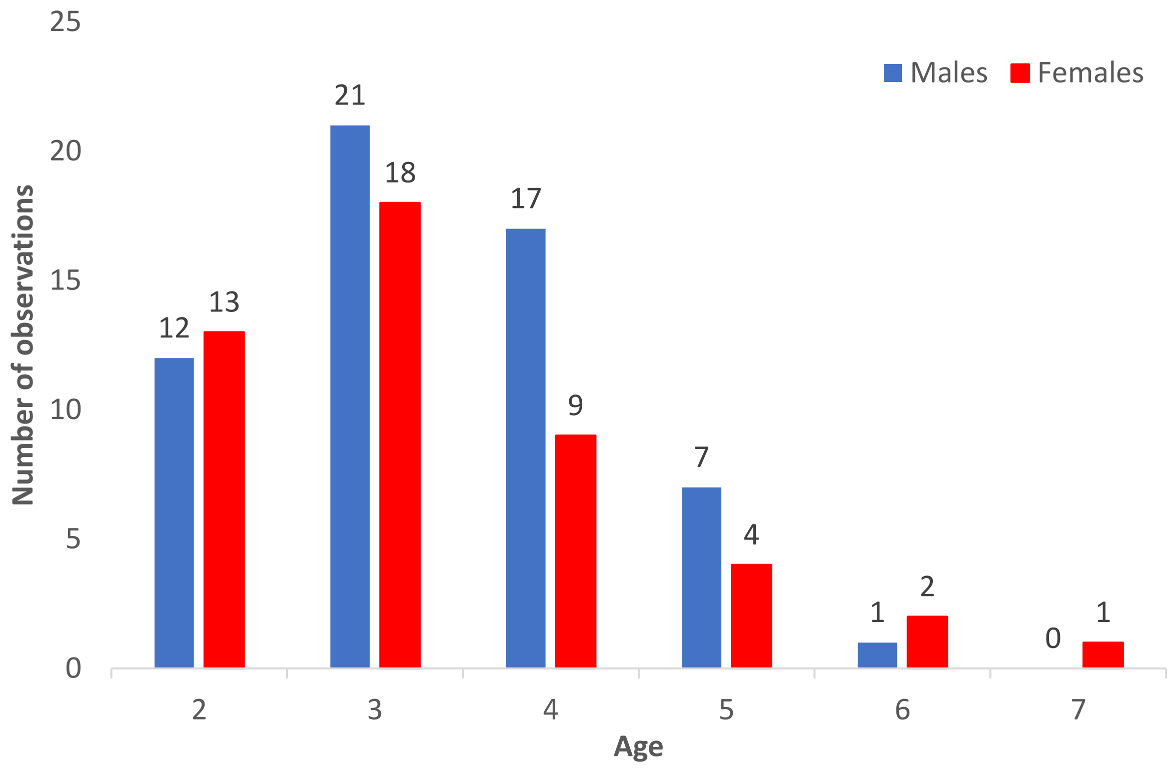

**Figure S2** Inter-annual within individual changes of plasma testosterone in repeatedly captured males (A) and females (B). Plasma T levels are shown as residuals from minimal adequate model 1 (Table 1) after controlling for technical factors possibly affecting plasma T measurements. The line crosslinks values from the same individual in different years. The minimal estimated age based on bird ringing is shown (males: $N_{ind}$ = 28, Nobs = 58; females: $N_{ind}$ = 21, Nobs = 47)

(A)

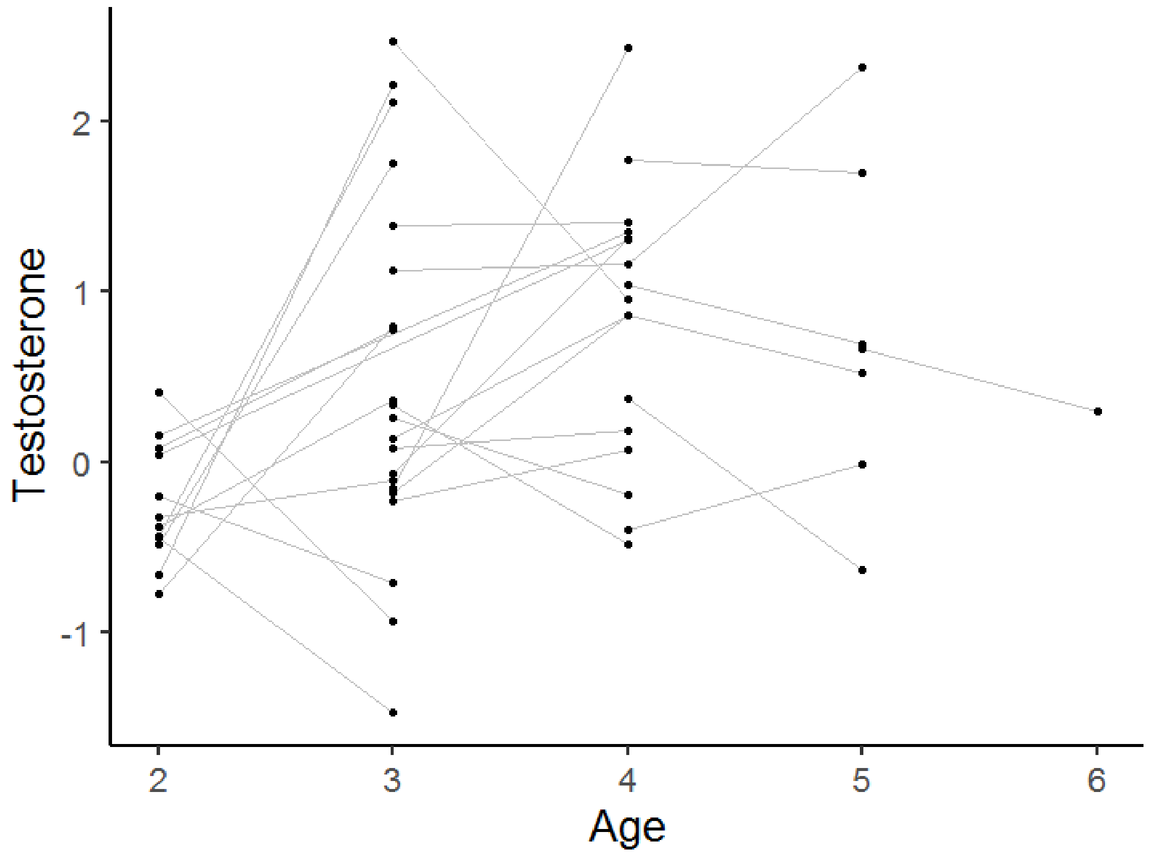

(B)

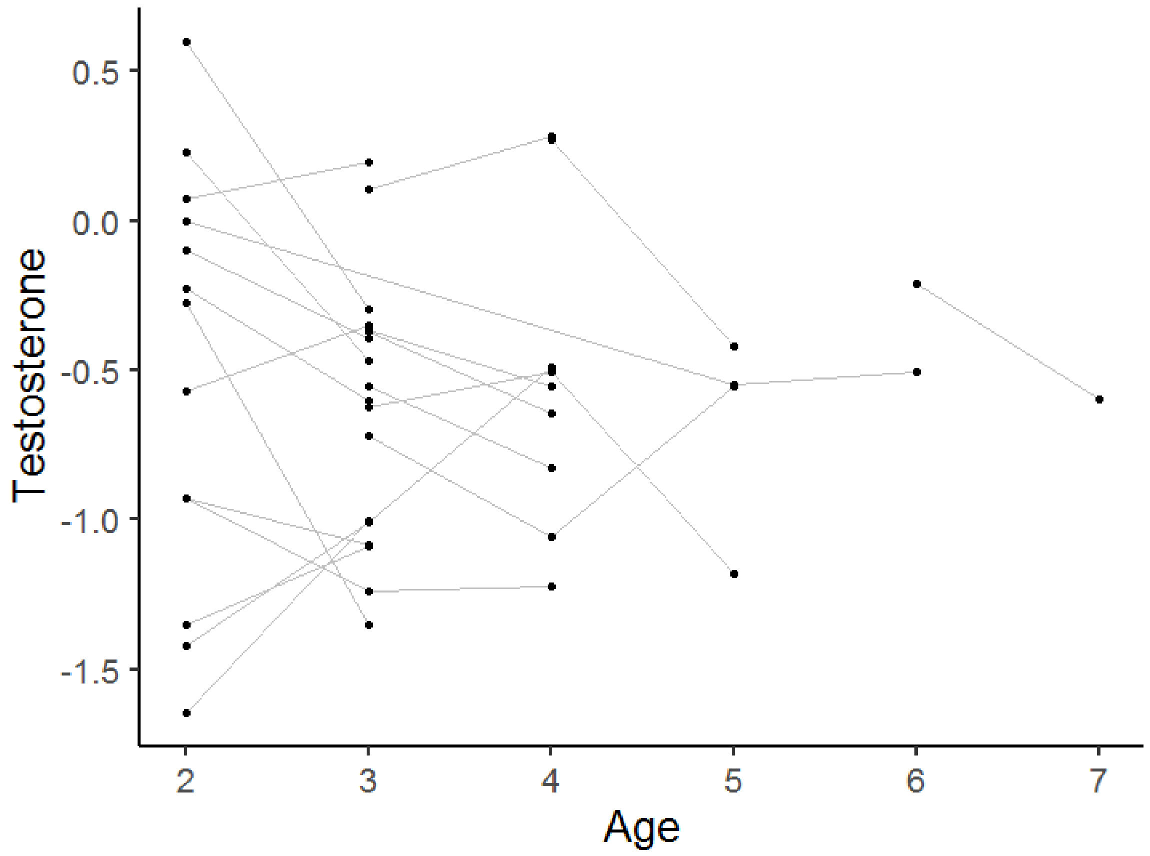

**Figure S3** Inter-annual within individual changes of breast stripe area in repeatedly captured males (A) and females (B). The line crosslinks values from the same individual in different years. The minimal estimated age based on bird ringing is shown (males: $N_{ind}$ = 28, Nobs = 58; females: $N_{ind}$ = 21, Nobs = 47)

(A)

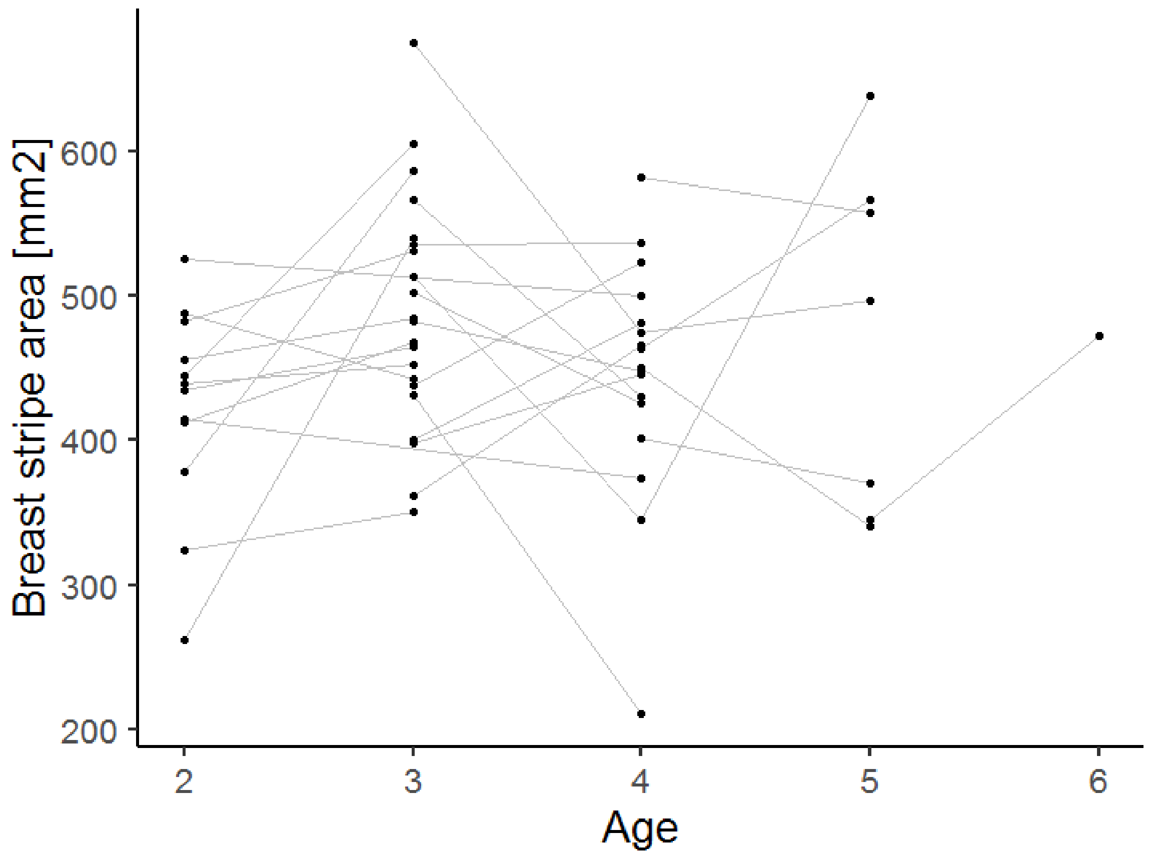

(B)

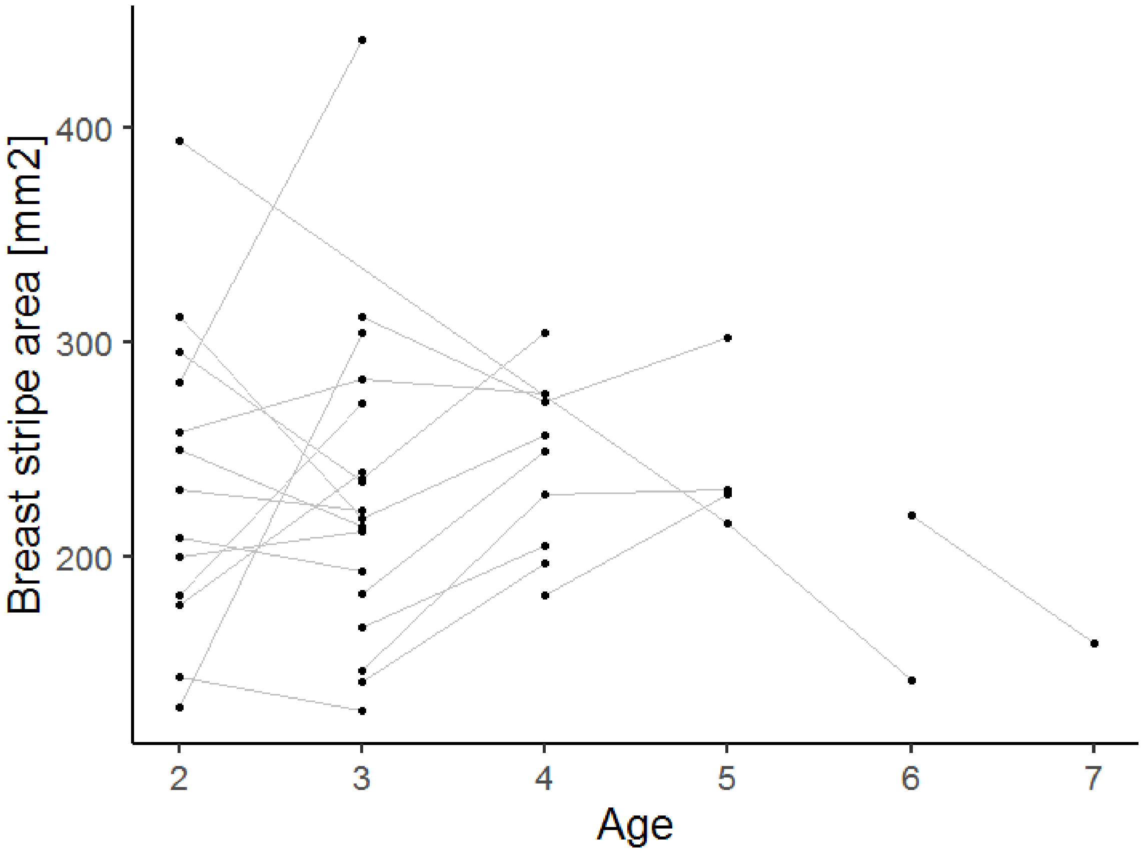

**Figure S4** Inter-annual within individual changes of yellow chroma in repeatedly captured males (A) and females (B). The line crosslinks values from the same individual in different years. The minimal estimated age based on bird ringing is shown (males: $N_{ind}$ = 28, Nobs = 58; females: $N_{ind}$ = 21, Nobs = 47)

(A)

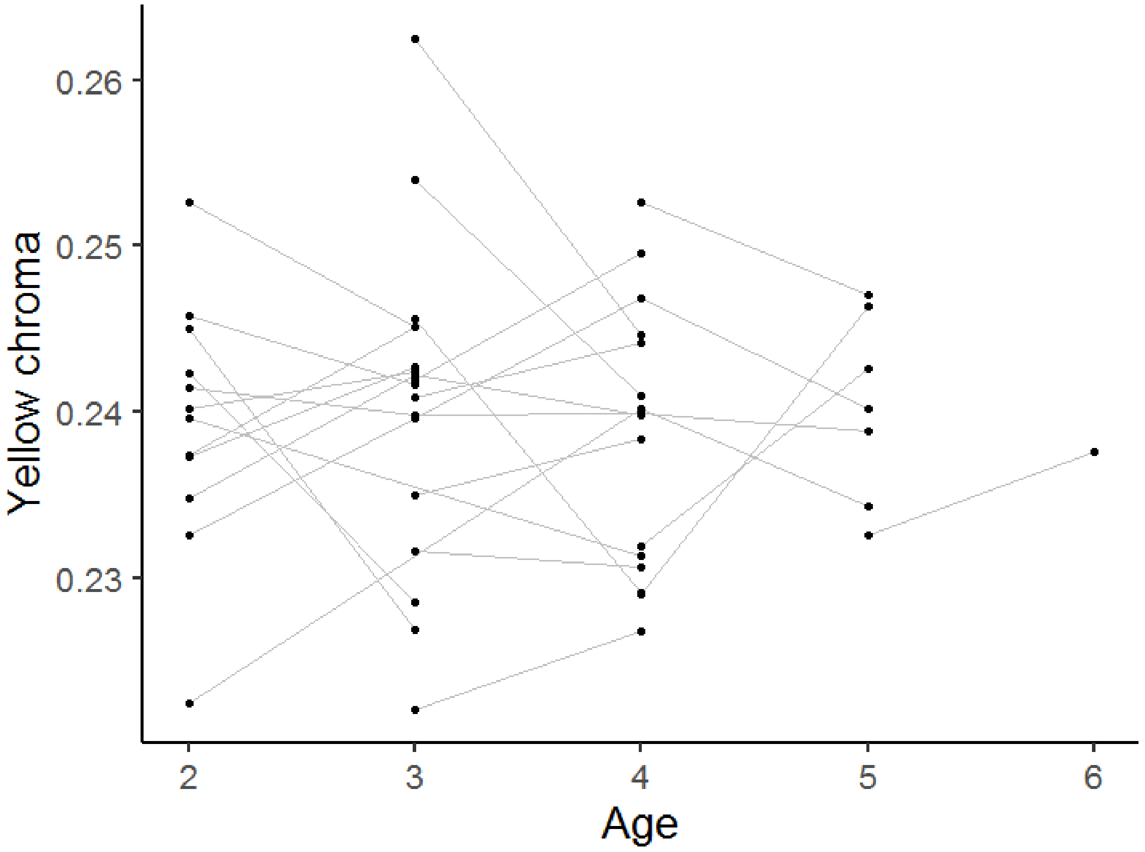

(B)

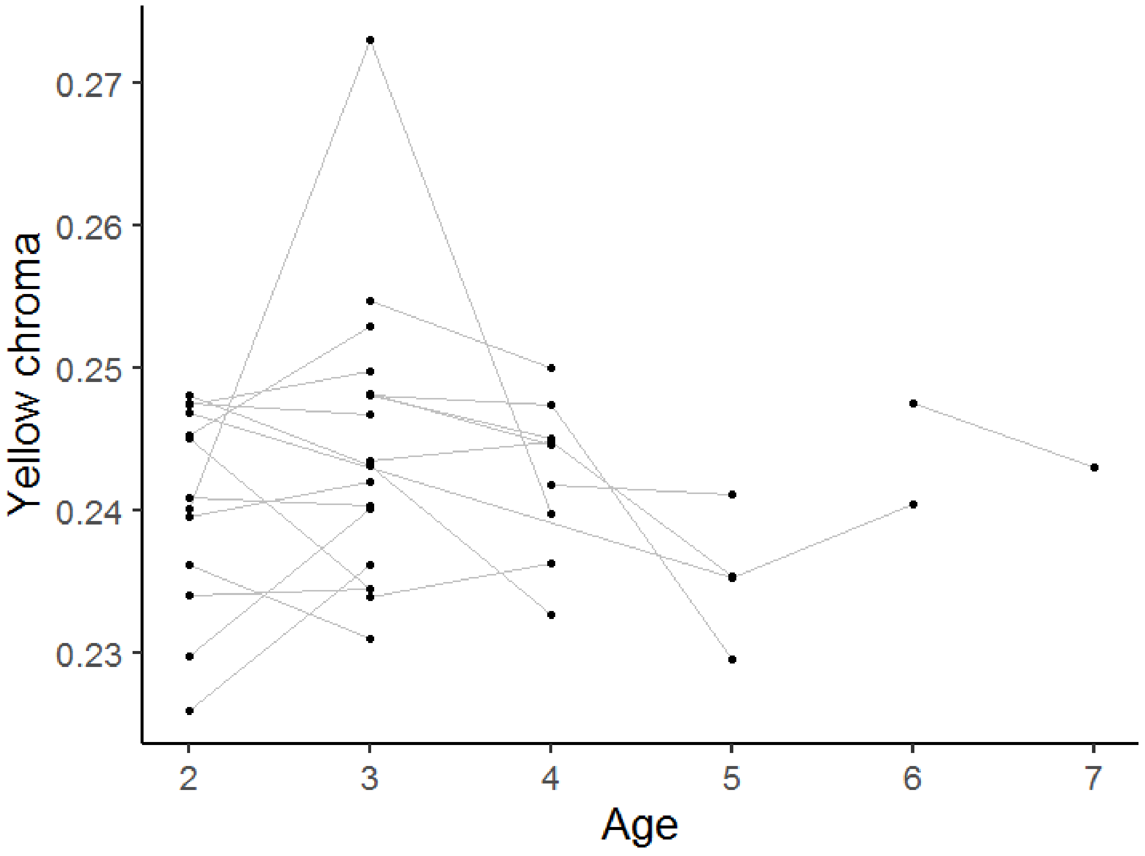

**Figure S5** Inter-annual within individual changes of yellow brightness in repeatedly captured males (A) and females (B). The line crosslinks values from the same individual in different years. The minimal estimated age based on bird ringing is shown (males: $N_{ind}$ = 28, Nobs = 58; females: $N_{ind}$ = 20, Nobs = 45)

(A)

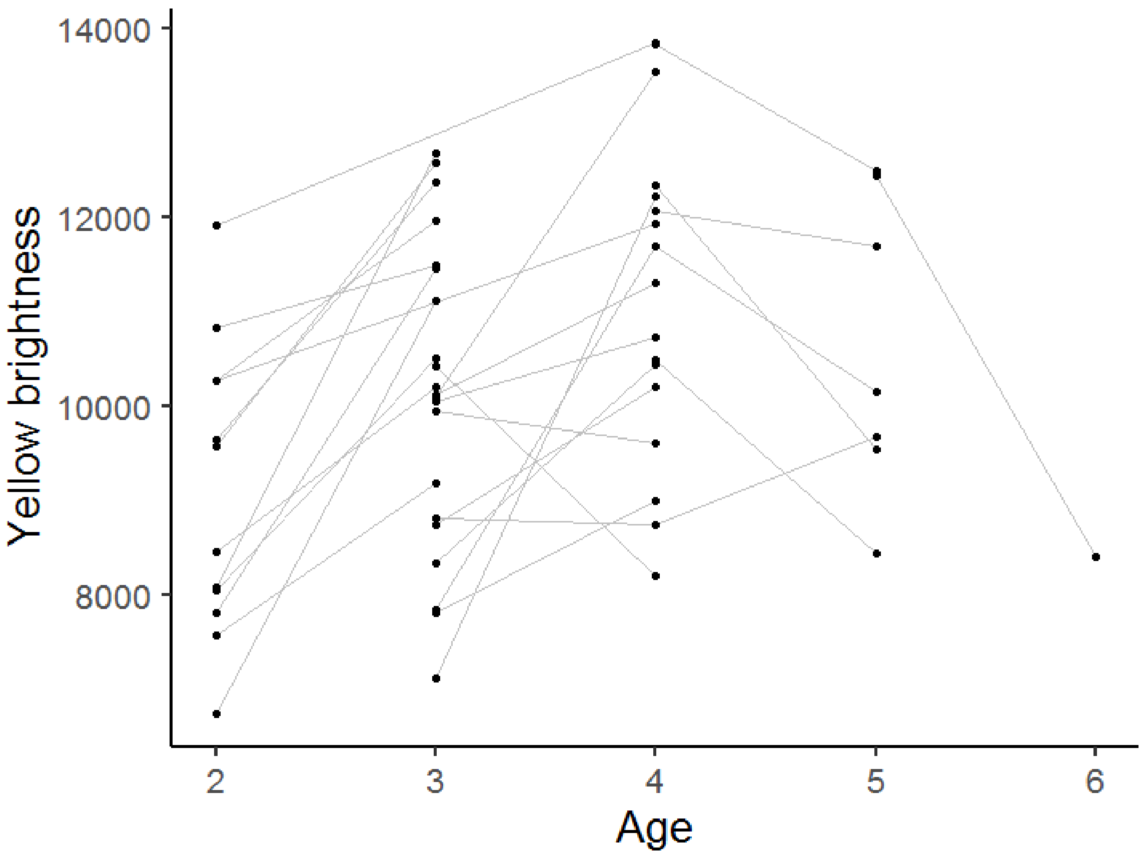

(B)

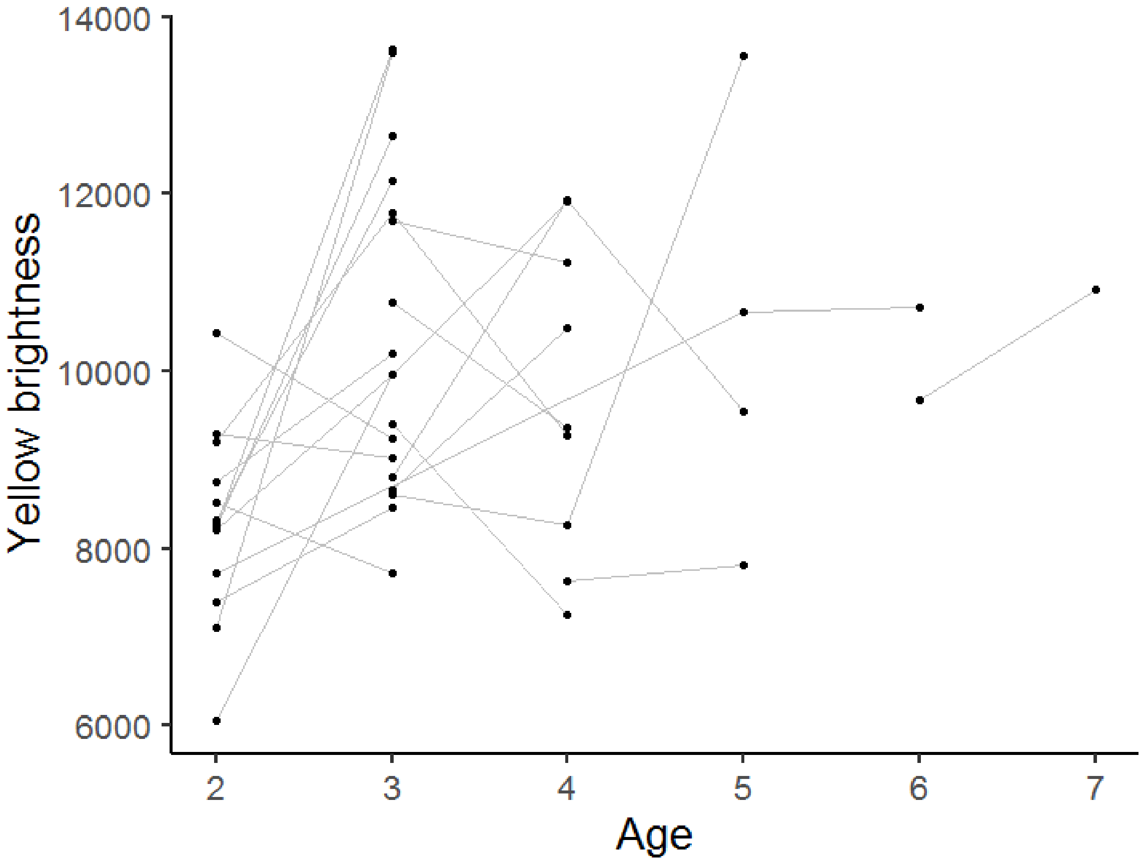

**Figure S6** Inter-annual within individual changes of feather growth rate (FGR) in repeatedly captured males (A) and females (B). The line crosslinks values from the same individual in different years. The minimal estimated age based on bird ringing is shown (males: $N_{ind}$ = 28, Nobs = 58; females: $N_{ind}$ = 21, Nobs = 47)

(A)

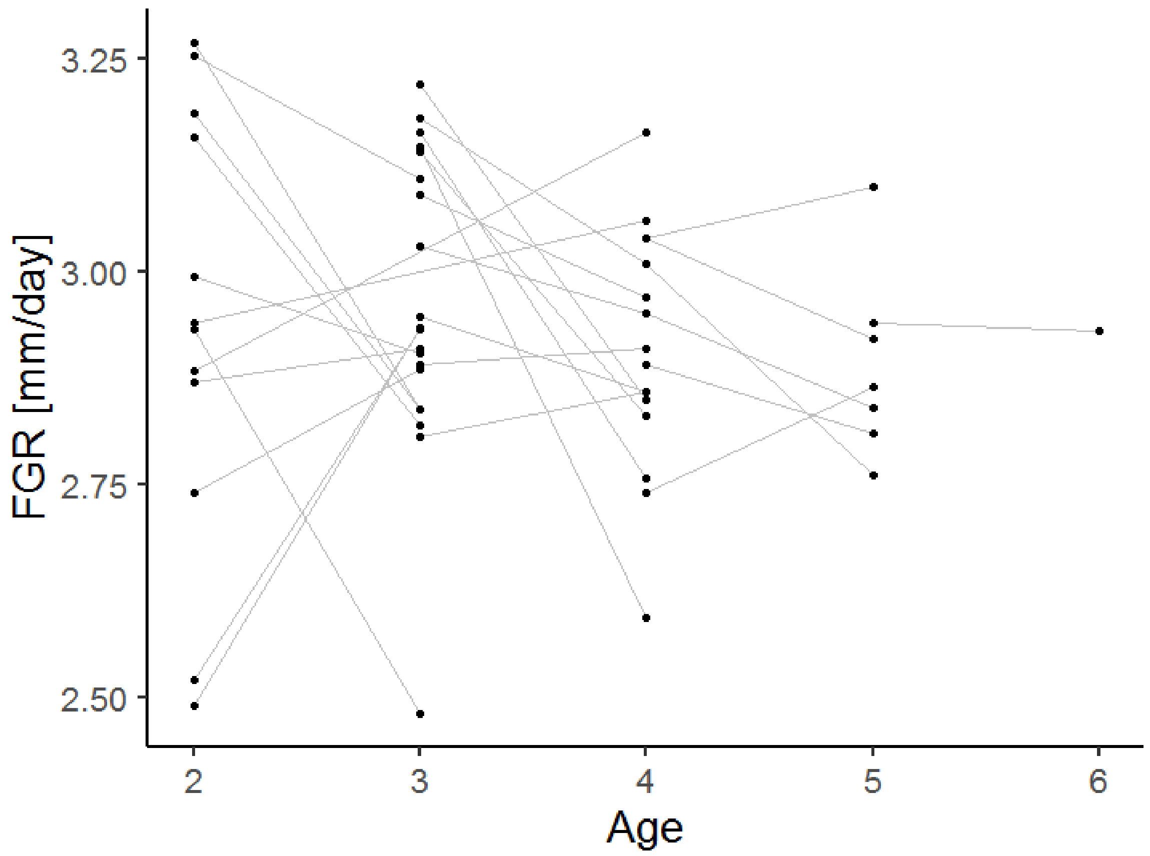

(B)

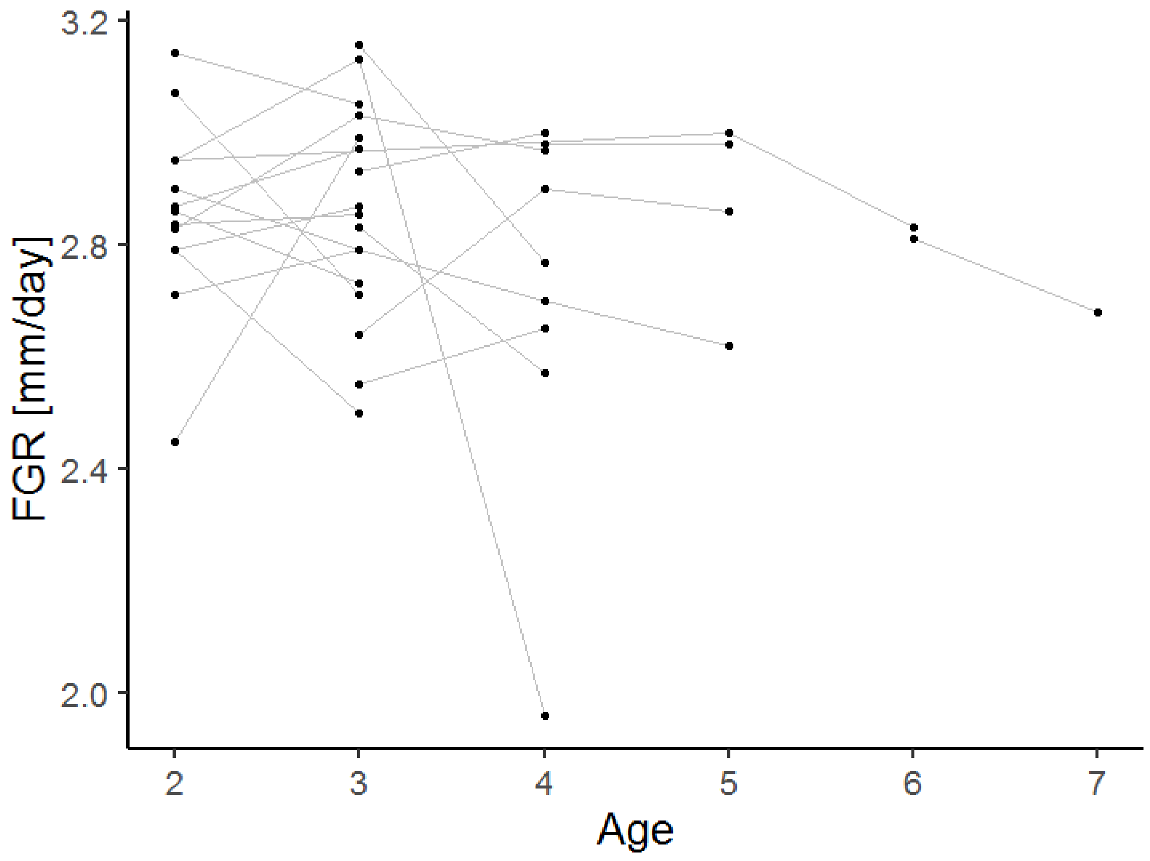

**Figure S7** Inter-annual within individual changes of heterophil: lymphocyte ratio (H/L ratio) in repeatedly captured males (A) and females (B). The line crosslinks values from the same individual in different years. The minimal estimated age based on bird ringing is shown (males: $N_{ind}$ = 28, Nobs = 58; females: $N_{ind}$ = 21, Nobs = 47)

(A)

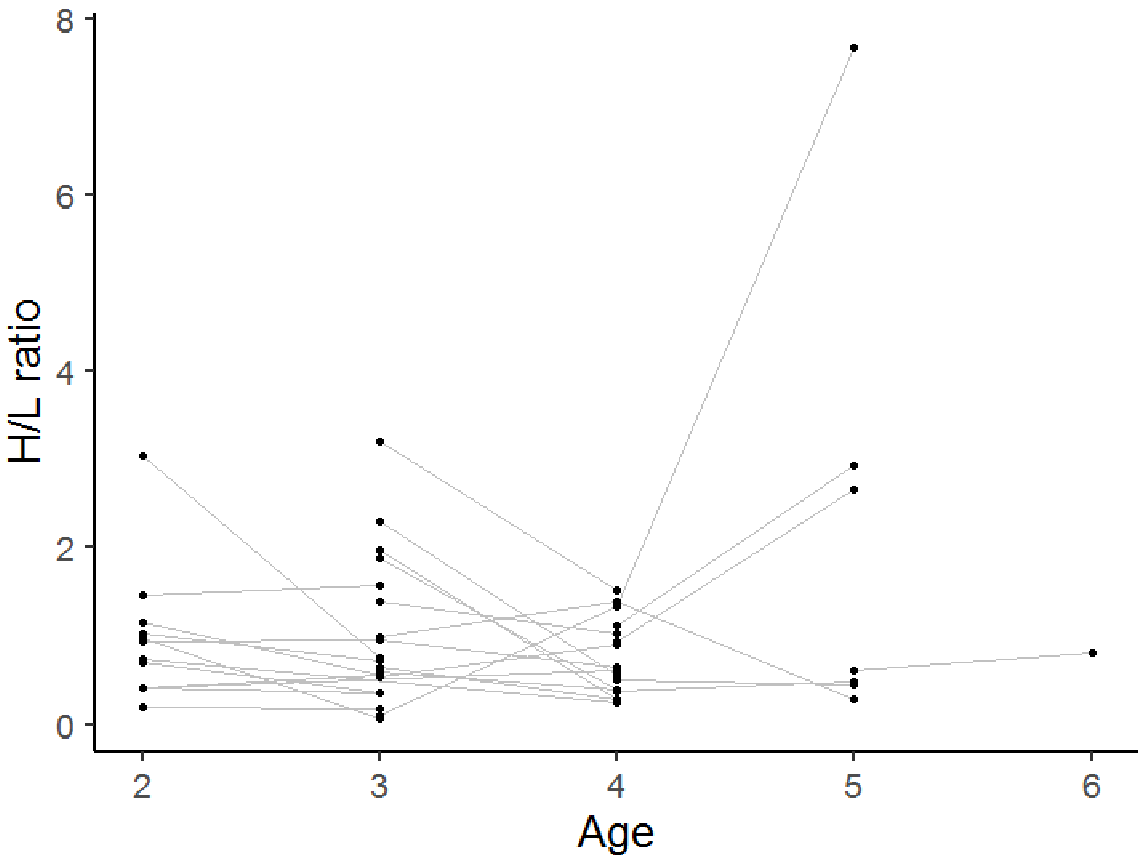

(B)

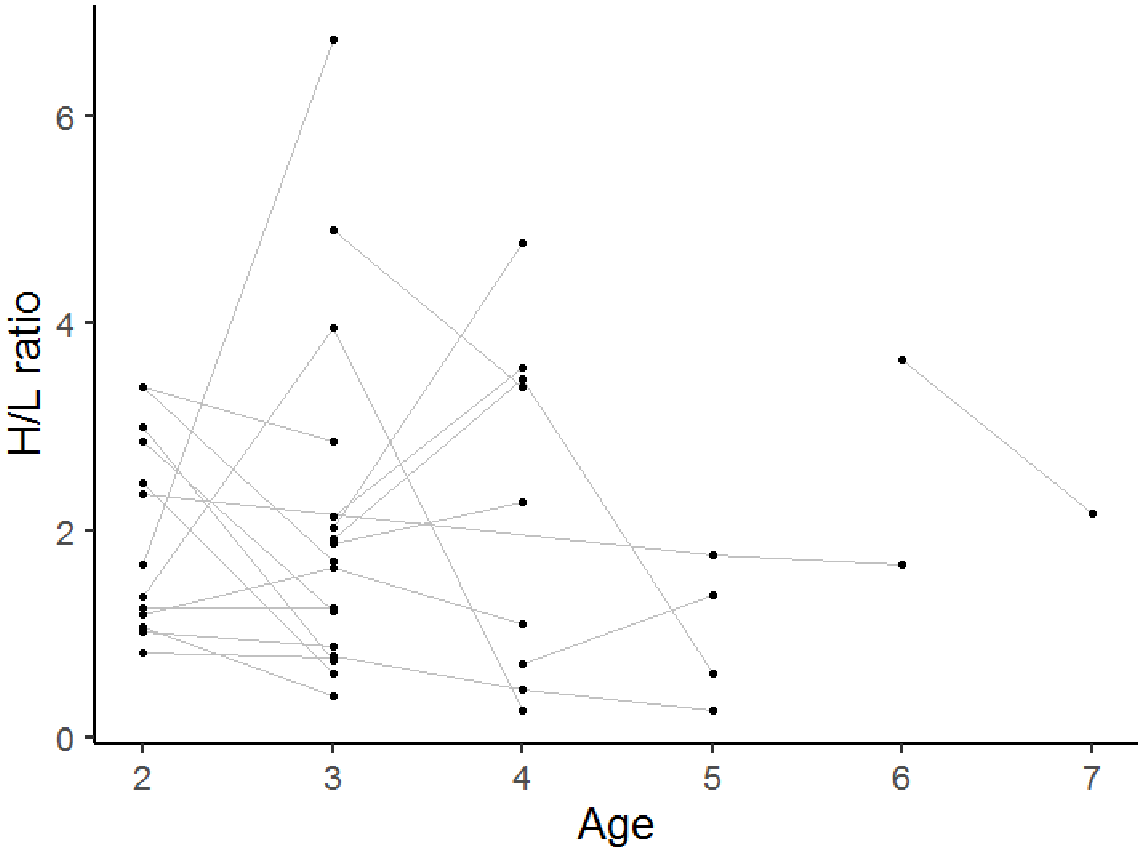

**Figure S8** Inter-annual within individual changes of size-standardised body mass (body mass) in repeatedly captured males (A) and females (B). The line crosslinks values from the same individual in different years. The minimal estimated age based on bird ringing is shown (males: $N_{ind}$ = 28, Nobs = 58; females: $N_{ind}$ = 21, Nobs = 47)

(A)

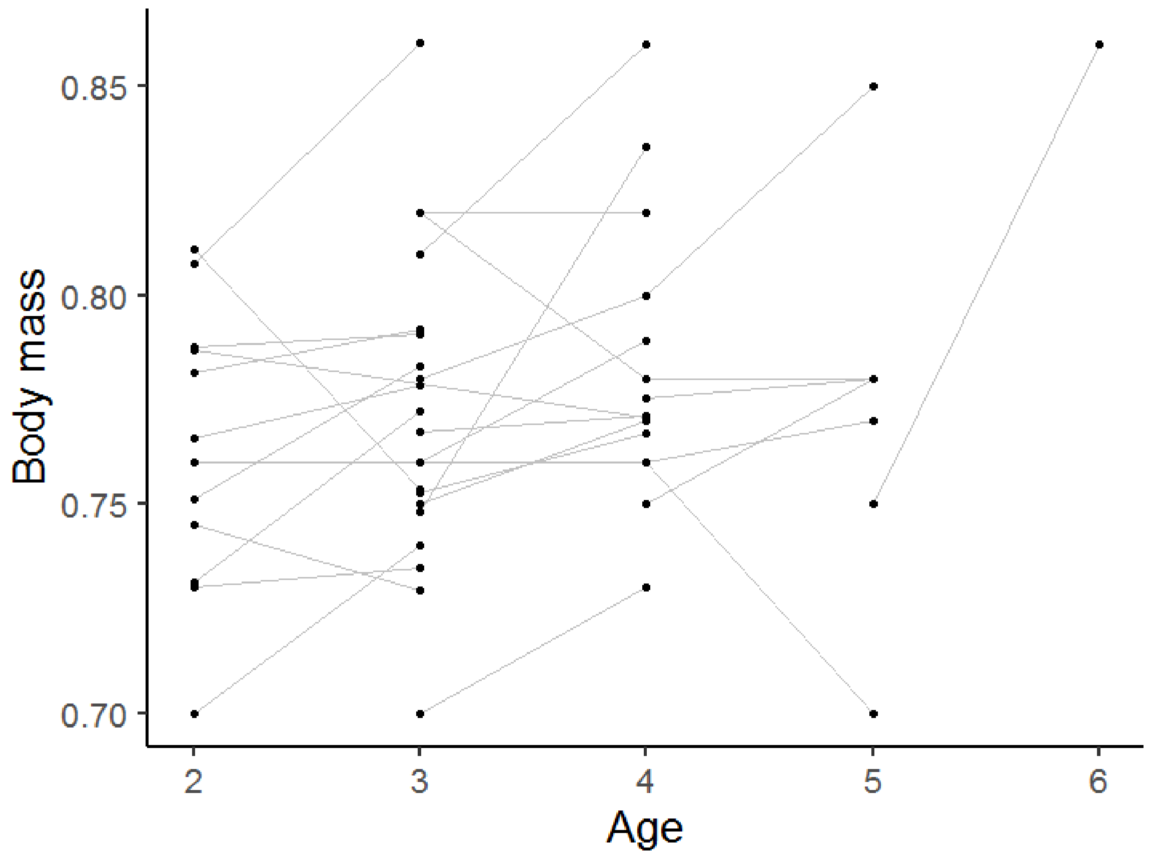

(B)

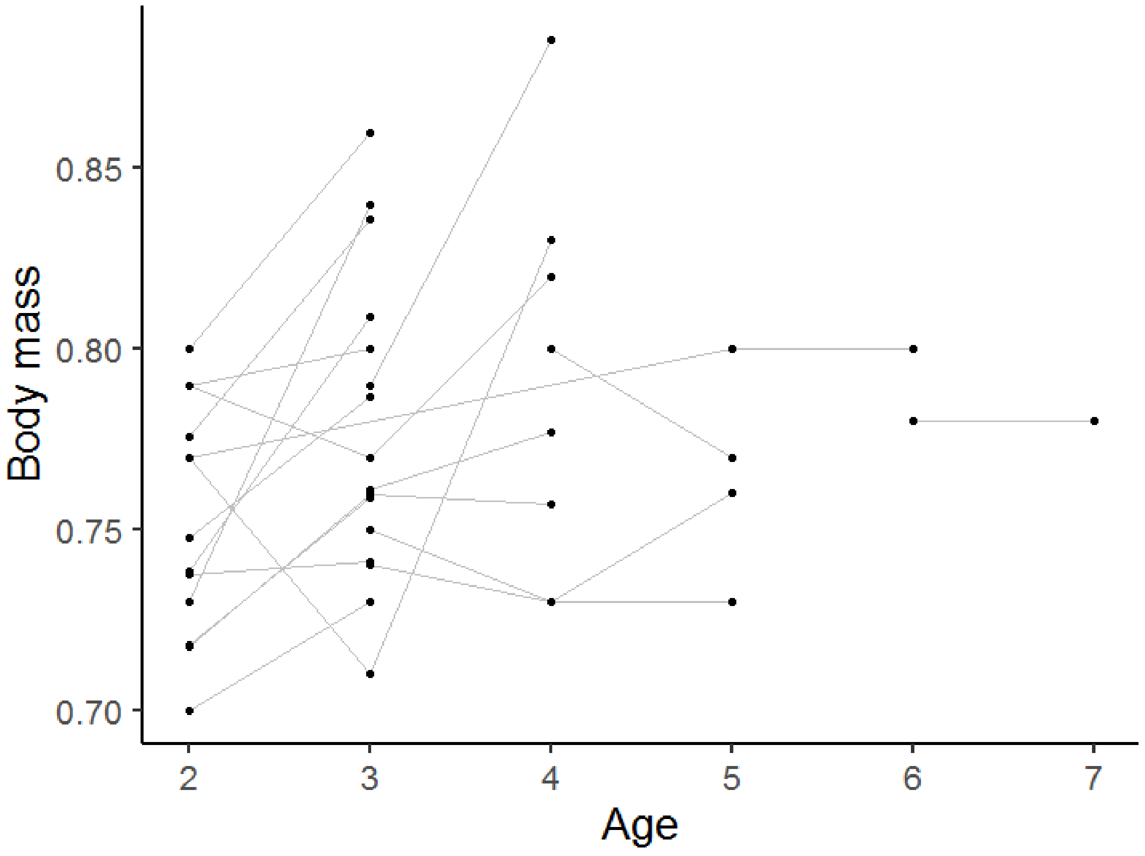

### References


Albrecht, T., Vinkler, M., Schnitzer, J., Polakova, R., Munclinger, P., & Bryja, J. (2009). Extra-pair fertilizations contribute to selection on secondary male ornamentation in a socially monogamous passerine. *Journal of Evolutionary Biology*, *22*(10), 2020–2030. doi: 10.1111/j.1420-9101.2009.01815.x

Vinkler, M., Schnitzer, J., Munclinger, P., Votýpka, J., & Albrecht, T. (2010). Haematological health assessment in a passerine with extremely high proportion of basophils in peripheral blood. *Journal of Ornithology*, *151*(4), 841–849. doi: 10.1007/s10336-010-0521-0
